# Binding of extracellular vesicles to stretched von Willebrand factor promotes platelet activation

**DOI:** 10.1101/2025.04.30.650971

**Authors:** Yuanyuan Wang, Xiaobo Liu, Tomasz Downar, Alper Topuz, Alexander T. Bauer, Katrin Nekipelov, Santra Brenna, Gerd Bendas, Berta Puig, Stefan W. Schneider, Dmitry A. Fedosov, Christian Gorzelanny

## Abstract

Von Willebrand factor (vWF), promoting platelet aggregation in various diseases such as COVID-19, malaria and cancer, is a huge multimeric glycoprotein. This extraordinary size makes vWF a unique shear stress sensing molecule. Below a critical shear stress, vWF is in a globular conformation that prevents platelet binding. Above the critical shear stress, vWF is stretched into platelet accessible fibers. Although previous studies have suggested that leukocytes or cancer cells can bind to vWF fibers, acting forces and the likelihood of cell adhesion has remained largely unexplored. Here, we report that vWF is a size-selective protein that prefers to interact with objects smaller than 4 μm in diameter. Consistently, tumor cell-derived extracellular vesicles (EVs) were able to interact with vWF in parallel to platelets. Although whole tumor cells under flow were unable to bind to vWF *per se*, binding of EVs and platelets along the vWF fiber promoted platelet aggregation, which in turn entrapped circulating tumor cells. In conclusion, our study highlights the shear-sensitive nature of vWF and its ability to bring EVs and platelets together to enhance coagulation. While EVs-vWF-platelet aggregates may serve as novel biomarkers, their therapeutic disruption may prevent hypercoagulation in disease.

## Introduction

The bloodstream hosts various components, among which von Willebrand Factor (vWF) plays a dominant role in coagulation and homeostasis. Synthesized in megakaryocytes and endothelial cells, ^1^ vWF represents the largest adhesive ligand in circulation. After synthesis, pro-vWF undergoes dimerization and multimerization, yielding large multimers that are either constitutively released or stored in the Weibel-Palade bodies of endothelial cells or the alpha-granules of platelets. ^2^

Upon activation of endothelial cells, typically triggered by various signals related to vascular stress, injury or inflammation, ^3^ vWF multimers stored within the endothelial cells are instantaneously released into the lumen of the blood vessel. This release plays a key role in promoting thrombosis by aiding platelet adhesion and aggregation at sites of endothelial cell activation. ^4^ The multimeric vWF is one of the largest protein assemblies within the mammalian body with sizes typically above 1×10^6^ kDa. In the Weibel-Palade bodies, vWF multimers are in a globular conformation. After their release from the endothelium and exposure to the blood flow vWF multimers unroll to form vWF fibers with a length of up to several hundred micrometers. ^5, 6^ The ability of vWF to unroll under physiological blood flow conditions is directly linked to the size of the multimer, which defines the target area of the shear force. ^7^ Previous numerical simulations suggest that the size of the vWF multimers is perfectly adjusted to the physiological shear forces acting in our vascular systems to achieve its essential function in primary hemostasis. ^8^

The conformational change of vWF multimers to elongated fibers opens the binding site for platelets within the A1 domain of vWF. Platelet binding to vWF is driven by strong electrostatic forces between the positively charged A1 domain and the negatively charged receptor on platelets, glycoprotein Ib alpha (GPIbα).

Extraordinary release of vWF from the endothelium under severe disease conditions such as sepsis, malaria or cancer increases the risk for intravascular thrombosis and the occlusion of microvessels. ^9–11^ The development of such pathophysiological situation is further supported by the lack of the vWF-cleaving enzyme ADAMTS13 (a disintegrin and metalloproteinase with a thrombospondin type 1 motif, member 13). ^12^ Beside the hereditary lack of ADAMTS13 due to gene mutations or the acquired deficiency due to the formation of autoantibodies, reduced plasma levels of ADAMTS13 are caused by its consumption at tissue sites with increased vWF secretion such as the tumor microenvironment. ^12^

Previous studies have suggested that vWF interacts not only with platelets but also with erythrocytes, ^10^ leukocytes^13^ and tumor cells through the A1 domain. ^1, 12, 14^ Related to this, the positive charge of the opened A1 domain suggests that in addition to GPIbα, other negatively charged biomolecules can interact with vWF. We have recently shown that vWF interacts with tumor cells through negatively charged heparan sulfate chains.^4^ Heparan sulfate (HS) is a linear sulfated glycosaminoglycan and belongs to the most negatively charged biopolymers, which is often expressed at cell surfaces in high abundance. ^15^ The anticoagulant heparin is a highly sulfated version of HS and synonymously used in the following. Although electrostatic interactions are strong and in principle sufficient to promote the binding of flowing cells to vWF fibers, previous studies have not yet considered the balance of shear stress-dependent vWF elongation required for adhesion and the shear stress-dependent drag force promoting cell detachment. ^16^

The drag force increases with the size of the bound cell, suggesting that the binding of larger objects such as tumor cells (diameter: **>** 10 μm) is less likely than the binding of platelets (diameter: 2-4 μm). In contrast, smaller objects such as extracellular vesicles (EVs), ranging from 0.05 μm to 5 μm, may have an increased ability to interact with vWF. EVs are released from various cell types, including platelets, leukocytes, and tumor cells. ^17^ They carry diverse cargos such as genetic materials, proteins, lipids, and small metabolites. ^18–20^ On their surface, EVs possess among others, HS-decorated proteoglycans and the prothrombotic tissue factor (TF). ^21, 22^ TF, the key initiator of the extrinsic coagulation cascade, acts in concert with coagulation factors VII and X, leading to thrombin generation and platelet activation. ^23^ Recent studies have further demonstrated that cancer-associated EVs drive thrombosis and metastasis via integrin β2-mediated platelet activation. ^24^ In line with that, clinical research suggested that EVs together with vWF are surrogate markers for hypercoagulation in diseases such as COVID-19, bacteremia and cancer. ^25, 26^ ^27^ While the contributions of EVs and vWF to thrombosis have been explored in previous research, investigations addressing the interaction between EVs and vWF, the contribution of shear stress, the influence of EV size, and the role of the A1 domain are missing.

In the present study, we aimed to systematically investigate the impact of shear stress on vWF elongation and the binding of flowing objects ranging from tumor cells and tumor cell-derived EVs to cell-like and EV-like particles. Experiments in microfluidic devices indicate that under physiological blood flow conditions vWF is size selective, which means that vWF prefers small objects, such as platelets and EVs. In contrast, the likelihood of larger object to be trapped by vWF such as whole tumor cells is very low. Lastly, we have shown that the binding of TF- expressing EV, in close proximity to platelets along the vWF fiber promoted platelet aggregation and activation, which in turn facilitated the entrapment of intact tumor cells under flow conditions.

## Materials and Methods

### 1. Cell lines and culture conditions

Human melanoma cell lines MV3, Mewo, and A2058 were cultured in RPMI-1640 medium (Sigma-Aldrich). The mouse melanoma cell line B16F10 was cultured in DMEM medium (Sigma-Aldrich). Both media were supplemented with 10% fetal bovine serum (FBS), 1% L-glutamine (Biochrom), and 1% penicillin/streptomycin (Biochrom).

Human umbilical vein endothelial cells (HUVECs) were isolated from human umbilical cords. The umbilical vein was cannulated and perfused with PBS to remove blood. Collagenase type I (0.1% w/v in PBS) was infused into the vein and incubated for 15-20 minutes at 37°C to digest the endothelial cells. The vein was then gently massaged to dislodge the cells. Isolated HUVECs were seeded into flasks pre-coated with 0.1% gelatin and cultured in Endothelial Cell Growth Medium-2 (EGM-2) (Sigma-Aldrich) supplemented with the provided growth factors, 10% FBS, and 1% penicillin/streptomycin. Only cells at passages 2-5 were used for experiments.

All cells were incubated in a humidified atmosphere at 37°C with 5% CO_2_ until they reached approximately 90% confluency.

### 2. HS coating of fluorescent polystyrene particles

Fluorescent polystyrene (PE) particles (0.5 μm PFP-0256, 1.8 μm PFH-2052, 4.0 μm PFP- 4070, 10.0 μm PFP-10052 from Kisker Biotech) were incubated with 1 mg/mL streptavidin in 0.1 M phosphate buffer (pH 7.4). The mixture was vortexed and incubated overnight at 4°C. After incubation, the suspension was centrifuged at 3000 × g for 15 minutes, and the supernatant was carefully removed. The pellet was washed with 4 mL isotonic buffered saline (IBS), centrifuged again, and the supernatant was removed. The pellet was then resuspended in IBS and mixed well to obtain a 5% w/v streptavidin-coated particle suspension.

Subsequently, 0.2 mL of the streptavidin-coated particle suspension was incubated with 2.0 mL of 100 μg/mL Biotin-Heparin (Sigma Aldrich) in 0.1 M sodium phosphate buffer (pH 5.5). The mixture was vortexed and incubated overnight at 4°C. After incubation, the suspension was centrifuged at 3000 × g for 10 minutes, and the pellet was washed three times with 4 mL of 0.1 M sodium phosphate buffer (pH 5.5). After each wash, the supernatant was carefully removed. Finally, the pellet was resuspended in 0.1 M sodium phosphate buffer (pH 5.5) to obtain a 1% w/v suspension.

### 3. EV isolation

EVs were isolated from cell culture medium using a commercial isolation kit (total exosome isolation kit, Invitrogen), following the manufacturer’s instructions. Briefly, cells were cultured in T75 flasks in 12 mL serum-free medium (Opti-MEM, Thermo Fischer) for 24 hours. The culture media was collected and centrifuged at 2000 × g for 30 minutes at 4°C to remove cells and debris. The supernatant was carefully transferred to a new tube, and 0.5 volumes of the total exosome isolation reagent (Invitrogen) were mixed and incubated at 4°C overnight. After incubation, the samples were centrifuged at 10,000 × g for 1 hour at 4°C. The supernatant was discarded, and the pellet containing the isolated EVs was resuspended in phosphate buffered saline (PBS) for further analysis.

### 4. Fluorescence labelling of EVs

After the isolation from cell culture medium, EVs were labelled with the lipophilic tracer DiD (1,1′-Dioctadecyl-3,3,3′,3′-tetramethylindocarbocyaninperchlorat, Thermo Fischer). The labelling was performed on ice and in the dark for 10 minutes. To remove any unincorporated dye, the labelled EVs were washed with PBS and centrifuged at 100,000 × g at 4°C for 60 minutes. The resulting pellet containing the labelled EVs was resuspended in PBS and used for the experimental procedures.

### 5. Single molecule force spectroscopy

Single molecule force spectroscopy (SMFS) measurements were performed with an atomic force microscope (AFM) (NanoWizard, JPK) as previously reported. ^1^ AFM tips (CS38/No Al, microMasch) and glass slides (Thermo Fisher Scientific) were functionalized with a polyethylene glycol (PEG) linker (Nanocs) and streptavidin was covalently attached to the linker through amino functionalization. Biotinylated HS (Sigma-Aldrich) was coupled to the streptavidin functionalized surfaces. Prior to the SMFS experiment, recombinant vWF in PBS (1μg/ml) was added to the glass slide and force distance cycles were immediately initiated. Rupture forces were recorded at different loading rates. At least 500 force curves per loading rate have been analyzed. The off rate and the bond length of the interaction between vWF and HS were determined as previously reported. ^28^

### 6. Surface acoustic wave biosensor measurements

Surface acoustic wave (SAW) measurements were conducted using a Sam®5 Blue acoustic biosensor (SAW Instruments GmbH) to analyze binding interactions between vWF and EVs. A model membrane containing 20 mol% 1,2-Di-(9Z-octadecenoyl)-sn-glycero-3-[(N-(5-amino- 1-carboxypentyl)iminodiacetic acid)succinyl nickel salt] (DGS-NTA(Ni)) and 80 mol% dipalmitoylphosphatidylcholine (DPPC, Sigma-Aldrich) was prepared on the sensor surface as previously described. ^29^ Baseline equilibration was achieved under buffer flow (40 μL/min). For vWF-EV interaction studies, 4.9 μg his-tagged vWF was injected at a flow rate of 20 μL/min to enable binding to the DGS-NTA(Ni) lipids in the membrane. The measurements were run at room temperature with a continuous buffer flow of 40 μL/min. After each injection of EVs, a rinse injection of 2 M NaCl solution was carried out in order to remove the EVs from the protein. A dilution series of B16F10 wild-type (WT) or Ext1⁻/⁻ EVs (1:200 to 1:25 in running buffer: 1 mM CaCl₂, 1 mM MgCl₂, 0.5 mM MnCl₂) was then injected, and real-time phase shifts were recorded.

### 7. Mouse experiments

All animal experiments were approved by the governmental animal care authorities (“Behörde für Justiz und Verbraucherschutz, Hamburg” project N033/2020). C57BL/6J mice and vWF deficient mice (Jackson Laboratory) were maintained under specific pathogen free conditions. Melanoma cells were injected intradermally (i.d.) into 8-12 week-old wild-type (WT) and von Willebrand factor knockout (vWF-/-) mice. After two weeks, HS-coated fluorescent nanocapsules (diameter: 157 ± 53 nm) were administered intravenously (i.v.). Twenty-four hours after nanocapsule administration, the mice were sacrificed. Tumor blood vessels containing accumulated nanocapsules were embedded for cryosectioning and subsequently analysed by immunofluorescence analysis.

### 8. EVs characterization Western blot

The total protein content in EVs or cell lysates were determined using the Micro BCA Protein assay kit (Thermo Scientific), following the manufacturer’s instructions. After quantification, the samples were denatured at 70°C for 10 minutes with NuPage LDS Sample Buffer (Invitrogen) and NuPage sample reducing agent (Invitrogen). Equal protein amounts were loaded onto NuPage 10% Bis-Tris gels (Invitrogen) and separated by electrophoresis. Proteins were transferred to a Transblot Turbo nitrocellulose membrane using wet-blotting. The membranes were then blocked for 1 hour and incubated with primary antibodies overnight at 4°C on a shaking platform. The following primary antibodies were used: rabbit antibody against Alix (1:400; #2171 CST, Millipore), rabbit monoclonal against CD81 (1:1000; #10037, Cell Signalling), mouse antibody against flotillin-1 (1:1000; #610820, BD Biosciences), and mouse antibody against GM130 (1:1000; #610822, BD Biosciences). After washing with TBST (Tris-buffered saline with Tween), the membranes were incubated for 1 hour with the respective HRP-conjugated secondary antibodies (1:1000, Cell Signaling) and subsequently washed six times with TBST. The protein bands were visualized using a chemiluminescence imaging system (azure biosystems, Dublin, USA).

#### Nanoparticle tracking analysis

EVs were subjected to Nanoparticle tracking analysis (NTA) for characterization, following a previously described protocol. ^20^ Briefly, 1 μL of the EVs suspension was diluted at 1:5000 in PBS and 500 μL of the diluted sample was loaded into the sample chamber of an LM10 unit (Nanosight). Ten videos of ten seconds were taken for each sample with a frame rate of 30 frames/sec. The movement of particles was subsequently analyzed using the NTA 3.0 software (Nanosight). The detection threshold was set to six and the screen gain to two.

#### Flow cytometry

The NovoCyte Quanteon flow cytometer was used to characterize EVs. Fluorescent polystyrene particles from Kisker Biotech with diameters of 0.1 μm (PFP-0245), 0.8 μm (PFP-0862), 2 μm (PFH-2052), and 4 μm (PFP-4070) served as size references to create a dimensional gate for EVs. Data analysis was performed using the Flowing Software (version 2.5.1).

#### Transmission electron microscopy

Transmission electron microscopy (TEM) was utilized as previously described. ^30^ Briefly, EV samples were prepared for negative staining. A carbon-coated formvar grid (2 nm carbon) was placed on a 20 μL drop of the EV sample, allowing it to adsorb for 10 seconds. The grid was subsequently washed three times with water drops and stained with a 3% uranyl acetate solution. After drying, the micrographs were captured using a transmission electron microscope (JEM 1400; JEOL Ltd) equipped with a bottom-mounted high-sensitivity 4K CMOS camera (TemCam F416; TVIPS).

#### Stimulated emission depletion microscopy

EVs were collected and labelled with DiD as described earlier. The labelled EV pellet were resuspended in 15 μL Fluoromount G (Southern Biotech). EVs were deposited on a glass slide and covered with a coverslip. Stimulated emission depletion (STED) microscopy experiments were conducted under similar conditions as previous studies. ^1^ Briefly, imaging was conducted in sequential line scanning mode with an Abberior STED expert line microscope. Excitation was achieved with a pulsed 640 nm laser, while depletion was performed using a near-infrared pulsed 775 nm laser. The dwell time for image capture was 3 ms, with a voxel size of 20 × 20 × 150 nm. Images were acquired in time-gating mode with a gating width of 8 ns and a delay of 781 ps. Data analysis was carried out by ImageJ software (version 1.54i).

### 9. Microfluidic experiment

#### EVs binding to endothelial vWF

HUVECs were cultured to confluence on fibronectin-coated μ-Slide I Luer (Ibidi) using Leibovitz’s L-15 Medium (Gibco) at 37°C. To stimulate the release of vWF, HUVECs were treated with 100 μM histamine under flow conditions.

To assess binding capacity, CellTrace™ calcein green-labeled (ThermoFisher) tumor cells (shown in blue in Figures) or DiD-labeled EVs (shown in black in Figures) were perfused through the BIOFLUX200 system in the presence of plasmatic vWF, with or without platelets or erythrocytes at a shear stress of 5 dyn/cm². Platelets, shown in green in the figures, were labeled by CellTrace™ calcein red-orange (ThermoFisher).

#### Binding of EVs or particles to recombinant vWF

BIOFLUX 200 48-WELL plates were coated with 20 μg/mL of recombinant wild-type vWF or recombinant vWF lacking the A1 domain (ΔA1 vWF) for 1 hour at room temperature. Tumor cell-derived EVs or HS-coated PE particles (0.5 μm PFP-0256, 1.8 μm PFH-2052, 4.0 μm PFP-4070, 10.0 μm PFP-10052) were perfused through the BIOFLUX200 system with or without platelets or erythrocytes. Applied shear forces ranging from 10 to 80 dyn/cm².

Adhesion of particles, platelets, tumor cells and EVs to vWF fibers was visualized by fluorescence microscopy (Observer Z.1, Zeiss). The resulting images were analyzed with the ImageJ software (version 1.54i) to quantify adhesion and binding events. Platelet coverage and aggregation were quantified based on fluorescence intensity.

### 10. Immunofluorescence staining of cells

Cells were fixed in 4% paraformaldehyde for 10 minutes, washed with PBS and blocked with 10% goat serum. The primary tissue factor antibody (Genetex SN20-16, 1:200) and secondary antibody: Alexa 555-conjugated goat anti-rabbit (IgG; Thermo Fisher Scientific, 1:500) were used for staining. Samples were imaged by fluorescence microscopy (Observer z.1, Zeiss). Image analysis was performed using the Image J software (version 1.52).

### 11. Flow cytometry for cells

Cells were collected and incubated with a TF antibody (rabbit anti human or mouse IgG, Gene Tex 1:100) for 1 hour on ice. After washing with PBS, TF antibodies were further labeled with Alexa Fluor 555 goat anti-rabbit IgG secondary antibodies (Thermo Fischer Scientific, 1:500) for 30 minutes on ice. Fluorescence signals were evaluated by flow cytometry (BD FACSCanto II, Biosciences). Data were analyzed by the Flowing Software (version 2.5.1).

### 12. Platelets and erythrocytes isolation

Citrated blood was centrifuged at 150 × g for 15 minutes without break, yielding three distinct phases: platelet-rich plasma with plasmatic proteins and platelets, the "buffy coat" interphase containing leukocytes, and the erythrocyte phase primarily composed of red blood cells. The platelet-rich plasma was gently transferred to a 15 mL conical bottom tube and mixed in a 1:1 ratio with washing buffer (136 mM NaCl, 2.7 mM KCl, 0.54 mM NaH_2_PO_4_ × H_2_O, 5 mM HEPES, 2.8 mM dextrose, pH 6.5), which included 1 U/ml of the platelet inhibitor apyrase. After washing, platelets were stained with CellTrace™ Calcein Red-Orange (ThermoFisher) for 15 minutes. Subsequently, the platelet was washed again and centrifuged at 1200 × g without break for 15 minutes. The supernatant was discarded, and the platelet pellet was resuspended in resuspension buffer (136 mM NaCl, 2.7 mM KCl, 0.54 mM NaH_2_PO_4_ × H_2_O, 11.9 mM NaHCO_3_, 5.6 mM glucose, 5% w/v bovine serum albumin (BSA), pH7.4). The erythrocyte pellet obtained from the citrated blood was transferred to a 50 mL tube and washed with an equal amount of HEPES-buffered Ringer’s solution (136 mM NaCl, 2.7 mM KCl, 1mM CaCl_2_, 1 mM MgCl_2_, 10 mM HEPES, 5 mM glucose) for three times. After each washing step, erythrocytes were centrifuged at 800 × g for 10 minutes. The supernatant was removed to obtain a purified pellet of washed erythrocytes.

### 13. Survival and migration of tumor cells

Tumor cell survival and migration were assessed using a basement membrane-like collagen matrix gel in μ-Slide 15 Well 3D ibiTreat chambers (ibidi). The gel was prepared using 100 μg/mL collagen I (Sigma-Aldrich) and precisely pipetted (10 μL) into the inner wells of the μ-Slide to achieve a meniscus-free, flat surface. Gelation was performed in a humidity chamber within an incubator for 30–60 minutes.

After gelation, 10,000 B16F10 cells, with or without platelets, were seeded onto the slides and incubated for 48 hours. The experiment was stopped by fixation of the gel with 4% paraformaldehyde. Gels were frozen and cut into 10 μm slices with a cryostat (CM3050S, Leica Biosystems). Nuclei were stained with DAPI; platelets were labeled with an antibody directed against thrombospondin 1 (Epredia MS-421-P). Fluorescence images were captured with a fluorescence microscope (Observer z.1, Zeiss). Aspect ratios were calculated as major/minor axis lengths using Image J (version 1.52).

### 14. Statistics

Statistical analysis was performed with Python (version 3.9.7) and GraphPad Prism 10 software and significance was tested by Student’s t-test and one-way ANOVA. All results are presented as means ± SD as indicated in the legend. P ≤ 0.05 was considered as significant difference.

## Results

### 1. Binding of platelets to vWF depends on shear stress

To investigate the impact of shear stress on the adhesive properties of vWF under flow, we applied microfluidic devices and live fluorescence microscopy. Prior to the experiment, platelets and erythrocytes were separated from whole blood and extensively washed to remove plasma proteins. A mixture of platelets (300,000/μL) and erythrocytes (45% hematokrit) was perfused over surfaces coated with recombinant vWF or, for control, BSA (**Figure 1A**). Platelets were stained with the fluorescence marker CellTrace^TM^ Calcein Red-Orange. The addition of erythrocytes enabled the margination of platelets towards the protein-coated surface at the bottom of the channel. ^8^ We performed force-ramp experiments, i.e. we increased the shear stress stepwise over 150 s from 10 dyn/cm² to 80 dyn/cm² (**Figure 1B**). The length of each step was 30 s. To change the shear stress, we increased the flow rate of the perfusing medium (**S-Figure 1**). Representative snapshots taken during the flow experiment are shown in **Figure 1C** and **Video S1**. The binding of platelets to vWF coated channels increased with increasing shear stress, whereas the binding of platelets to BSA-coated surfaces was comparable low and unchanged over the applied range of shear stresses. On vWF-coated channels, platelets adhered in a “pearl-on-string” fashion, consistent with previous reports and indicative of the elongation of the globular vWF to its stretched conformation and thus the opening of the platelet binding site within the A1 domain of vWF. ^31^ The quantitative evaluation of the flow experiment is shown in **Figure 1D**. The x-axis indicates the time of the experiment and the distinct force steps (arrows). Platelet adhesion on vWF-coated surfaces reaches the maximum after 140 s during the 80 dyn/cm² step and declines slightly afterwards suggesting that the applied drag force exceeds the adhesion force. Platelet adhesion to BSA-coated surfaces remained at a very low level over the course of the experiment. To better extract the impact of shear stress independent of the time interval, we calculated the binding rate (slope of platelet adhesion per shear stress segment). **Figure 1E** presents the binding rate as a function of the applied shear stress and shows a steep increase of platelet adhesion starting at 20 dyn/cm². The binding rate peaks at 60 dyn/cm² and declines towards 80 dyn/cm². As schematically summarized in **Figure 1F**, we concluded that in our experimental setup, vWF changed from a globular conformation at 10 dyn/cm² to an extensively stretched conformation at 60 dyn/cm² offering binding sites (open A1 domain) to platelets. At 80 dyn/cm², bound platelets or large platelet-vWF conglomerates (not shown in the schematic drawing) detach as the drag force pulling on the platelets exceeds the adhesion force.

**Figure 1.**
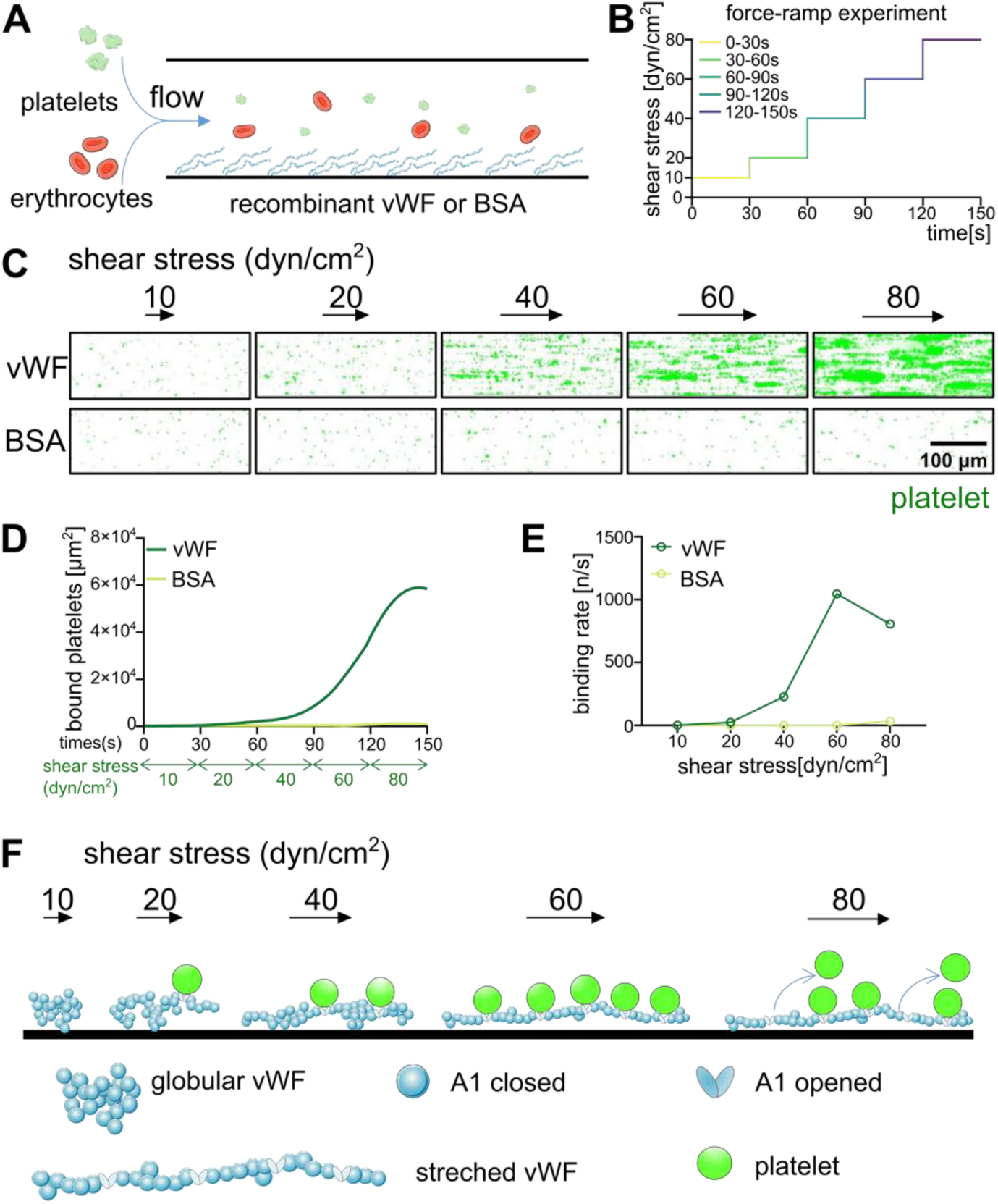
Binding of platelets to stretched vWF under shear stress. (**A**) Schematic representation of the microfluidic experiment. Platelets and erythrocytes (45% haematocrit) were perfused over microfluidic channels coated with recombinant vWF or BSA. (**B**) Force-ramp experiments were performed with a time-dependent change of the shear stress. (**C**) Time-lapse dual-color fluorescence images of the dynamic interaction of platelets (green) with vWF or BSA over the indicated shear stress range. (**D**) Binding of platelets (green) to vWF at increasing shear stresses over time. (**E**) Binding rates of platelets as a function of the applied shear stress. Data have been derived from the data shown in (D). (**F**) Schematic drawing showing the stretching of globular vWF under increasing shear stresses, the shear stress-dependent opening of the A1 domain and the binding of platelets to the open A1 domain. (Schematic elements used from Servier Medical Art: https://smart.servier.com/).

### 2. Stretched vWF favours the binding of small particles

This study intends to understand the effect of the size of flowing objects and their interaction with vWF at different shear stress conditions. For a standardized investigation, PE-particles with diameters of 0.5, 1.8, 4 or 10 μm were perfused over vWF-coated surfaces together with erythrocytes (**Figure 2A**). We compared microfluidic channels coated either with WT vWF or vWF lacking the A1 domain (ΔA1 vWF). In the previous experiment (**Figure 1)**, the time interval of each force step was 30 s independent of the flow rate. Consequently, more liquid is pumped over the surface at a higher than at a lower shear stress. To better compare the binding of the PE-particles, we adjusted the time interval of our force-ramp experiment. Therefore, the volume of medium and thus the number of particles pumped over the channel is identical in each force step (**Figure 2B**). To increase the affinity of the PE-particles to the A1 domain of vWF, we coated the particles with HS, which is a known ligand for the A1 domain and ubiquitously expressed on mammalian cells^1^ (**S-Figure 2**). SAW experiments confirmed the binding of HS to WT vWF compared to ΔA1 vWF (**S-Figure 2A**). After HS coating, PE-particles exhibited significantly increased binding to WT vWF (**S-Figure 2B–D**). **Figure 2C** shows representative snapshots of the flow experiment performed with the different particles, simultaneously. The number of the small particles (diameters: 0.5 μm and 1.8 μm) increased with increasing shear stress. The medium large particles (diameter: 4 μm) bound weakly above 40 dyn/cm² shear stress. The largest particles (diameter: 10 μm) did not bind at any of the tested shear stresses. The quantitative evaluation of the particle binding is shown in **Figure 2D**. Except for the 10 μm particles, the number of bound particles increases with time and increasing shear stress. The change of the shear stress is indicated in the figure by the color gradient of the curves (light colors correspond to lower shear stress; strong colors to higher shear stress). Consistent with the snapshots shown in **Figure 2C**, the smallest particles (diameter: 0.5 μm and 1.8 μm) bound best. To summarize multiple experiments (n=3) we calculated the binding rates for the tested shear stress range as shown in **Figure 2E**. The binding rates of the 0.5 and 1.8 μm particles increased until the final shear stress of 80 dyn/cm² was reached. The shear stress-dependent binding suggests that the elongation of vWF and thus the exposure of the A1 domain is required. In the next experiments, we measured the binding of particles to surfaces coated with the ΔA1 vWF. **Figure 2F** indicates that lack of the A1 domain abolished the binding of the particles and no shear stress related change of the binding affinity was measured (**Figure 2G**).

**Figure 2.**
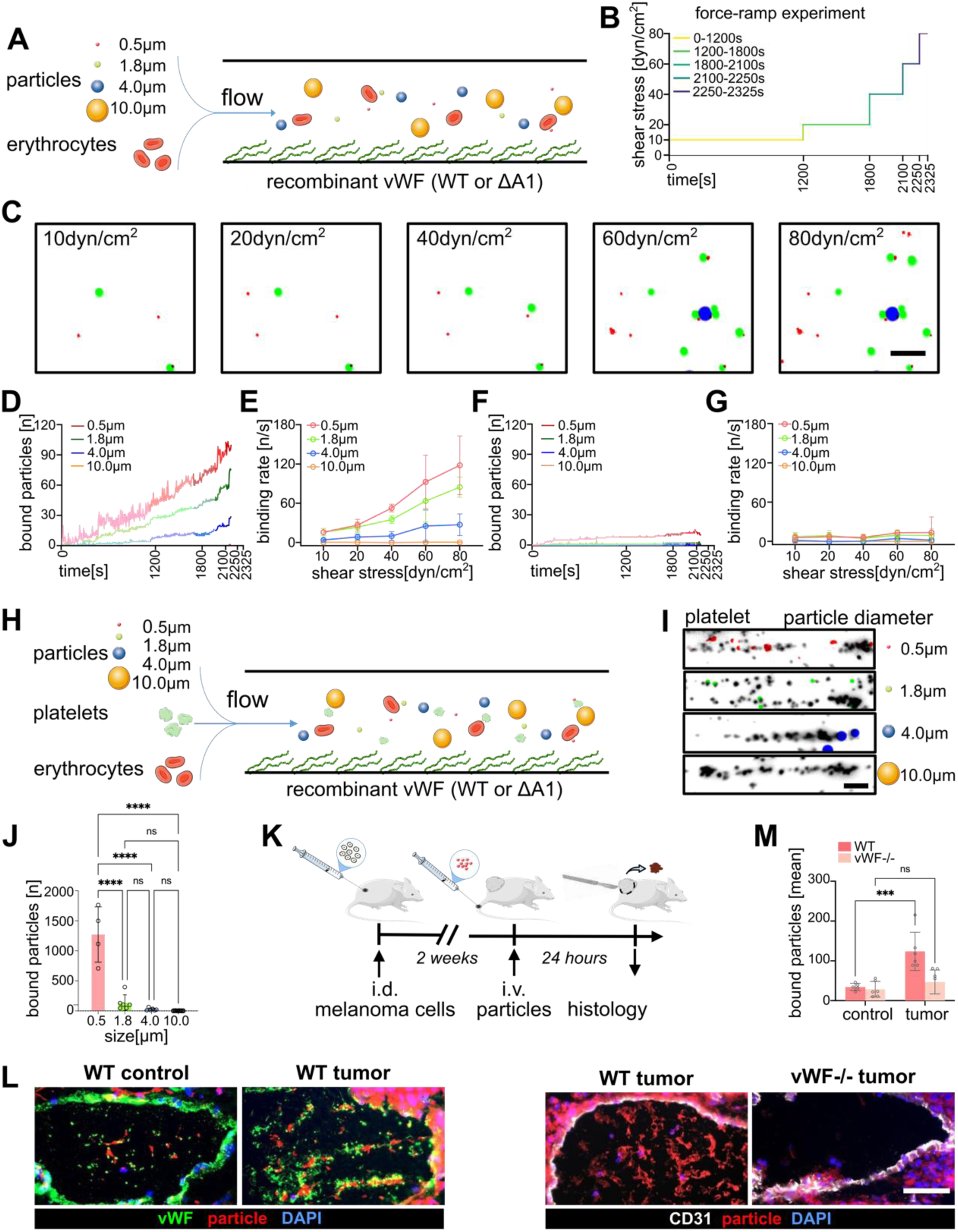
Binding of particles to stretched vWF under flow conditions. (**A**) Schematic representation of the microfluidic experiment. Particles of different sizes were perfused through vWF-coated channels in the presence of erythrocytes. (**B**) Force-ramp experiments were performed with a time-dependent change of the shear stress: 10 dyn/cm^2^ (0-1200 s), 20 dyn/cm^2^ (1201-1800 s), 30 dyn/cm^2^ (1801-2100 s), 40 dyn/cm^2^ (2101-2250 s), 60 dyn/cm^2^ (2251-2325 s), 80 dyn/cm^2^ (2326-2364 s). (**C**) Representative images of PE particles bound to WT vWF at the indicated shear stress. Scale bar = 10 μm. (**D**) Particle binding to WT vWF coated surfaces as a function of time and shear stress. The change of the shear stress is indicated by the color gradient (light colors: low shear stress; strong colors: high shear stress). (**E**) Binding rates of particles to WT vWF coated surfaces as a function of shear stress. (**F**) Particle binding to ΔA1 vWF coated surfaces as a function of time and shear stress. The change of the shear stress is indicated by the color gradient (light colors: low shear stress; strong colors: high shear stress). (**G**) Binding rates of particles to ΔA1 vWF coated surfaces as a function of shear stress. (**H**) Schematic representation of the microfluidic experiment. Particles with the indicated sizes were perfused through vWF-coated channels in the presence of platelets and erythrocytes. Shear stress was constantly at 60 dyn/cm². (**I**) Particles with diameters of 0.5, 1.8 and 4.0 μm bound to the channel surface in close proximity to platelets (gray). Particles with a diameter of 10μm did not bind. Scale bar = 10μm. (**J**) Quantitative evaluation of fluorescence particle binding in the presence of platelets (n = 4). (**K**) Schematic presentation of the mouse experiment. Melanoma cells were intradermally (i.d.) injected into wild-type (WT) and vWF knockout (vWF-/-) mice. After two weeks, DiD-labeled nanoparticles (diameter: 157 ± 53nm) were intravenously (i.v.) administered. Twenty-four hours later, the mice were sacrificed, and tumors were analyzed by immunofluorescence staining of the tissue. (**L**) In WT mice, nanoparticles (red) accumulated in the vasculature of tumors in co-localization with vWF (green). Lack of vWF in the control vessel of the skin prevented nanoparticle accumulation. In vWF-/- mice, nanoparticle accumulation was abolished. In vWF-/- mice the endothelial cell marker CD31 (white) was used to stain the vascular wall. Nuclei are stained by DAPI (blue). Scale bar = 100 μm. (**M**) Quantification of bound particles in the blood vessels of WT and vWF-/- mice, and control and tumor tissues. (n= 6). *** P ≤ 0.001, **** P ≤ 0.0001 one-way ANOVA with a Tukey post hoc test. (Schematic elements used from Servier Medical Art: https://smart.servier.com/).

To confirm our findings and to understand the difference in platelet and particle binding, we performed experiments with platelets together with the PE-particles (**Figure 2H**). The data shown in **Figure 2I**, indicate that compared to the particles, more platelet bound to the vWF-coated surface and that platelets tend to form aggregates. Compared to the experiments without platelets (**Figure 2D**), binding of the different particles to vWF was similar showing highest adhesion for the 0.5 μm articles and no binding of the 10 μm particles (**Figure 2J**).

In further experiments, we intended to demonstrate that the particle binding ability of vWF is of physiological relevance. Therefore, we injected non-toxic nanoparticles loaded with a fluorescence dye intravenously into tumor bearing mice (**Figure 2K**). The nanoparticles had a diameter of about 200 nm and were thus in the size-range accepted by vWF^32^. From our previous studies we knew that the tumor endothelium releases vWF into the blood vessel lumen, while the level of ADAMTS13 activity is decreased. ^12^ Consequently, vWF fibers are persistently formed. ^12^ In line with our previous findings, nanoparticles accumulated along the vWF fibers in the lumen of tumor blood vessels (**Figure 2L**), while vWF fibers were absent in healthy skin (control). Therefore, no nanoparticles accumulated in the lumen of control blood vessels. The lack of vWF in vWF knockout mice prevented nanoparticle accumulations in tumor blood vessels. **Figure 2M** shows the quantitative evaluation of the fluorescence images and indicates a significant relationship between vWF fiber formation in the blood vessel lumen and nanoparticle binding.

### 3. Theoretical calculations of particle binding to vWF-coated surfaces

To confirm our data with an independent approach, we performed theoretical calculations of the particle binding to vWF under shear stress. The calculation, described in detail in the Data Supplement and **S-Figures 3** and **S-Figure 4**, considers the balance between the adhesive forces (*F_ad_*) of a particle close to the vWF-coated surface and the drag force (*F_drag_*) pulling on the particle (**Figure 3A**). In our calculation we assumed that at a critical separation (*s_crit_*) of the particle from the channel surface *F_drag_* and *F_ad_* are at equilibrium. Particles closer to the surface will attach, whereas particles above *s_crit_* are not able to bind (**Figure 3B**). *F_drag_* is determined by the shear stress and the particle diameter. Therefore, we studied the effect of different shear stress conditions and the impact of the particle diameter. To allow the comparison to the experiments shown in **Figure 2**, we assumed the previous force ramp profile (**Figure 2B**). In **Figure 3C**, we show the increased binding of particles over time. Consistent with our experiments (**Figure 2)**, small particles bound best, while the largest particles (10.0 μm diameter) failed to bind to the surface. At lower shear stress (<40 dyn/cm²) the binding rate of the 0.5 μm and 1.8 μm particles increased with the increasing shear stress indicating that *F_ad_* is larger than *F_drag_* (**Figure 3D**). Above 60 dyn/cm², a plateau is reached indicating that at high shear stress *F_drag_* and *F_ad_* are balanced. Although some particles with a diameter of 4 μm could bind to the surface (**Figure 3C**), the effect of shear stress was neglectable (**Figure 3D**). We repeated the calculation assuming that the surface was coated with ΔA1 vWF (**Figure 3E, F**).

**Figure 3.**
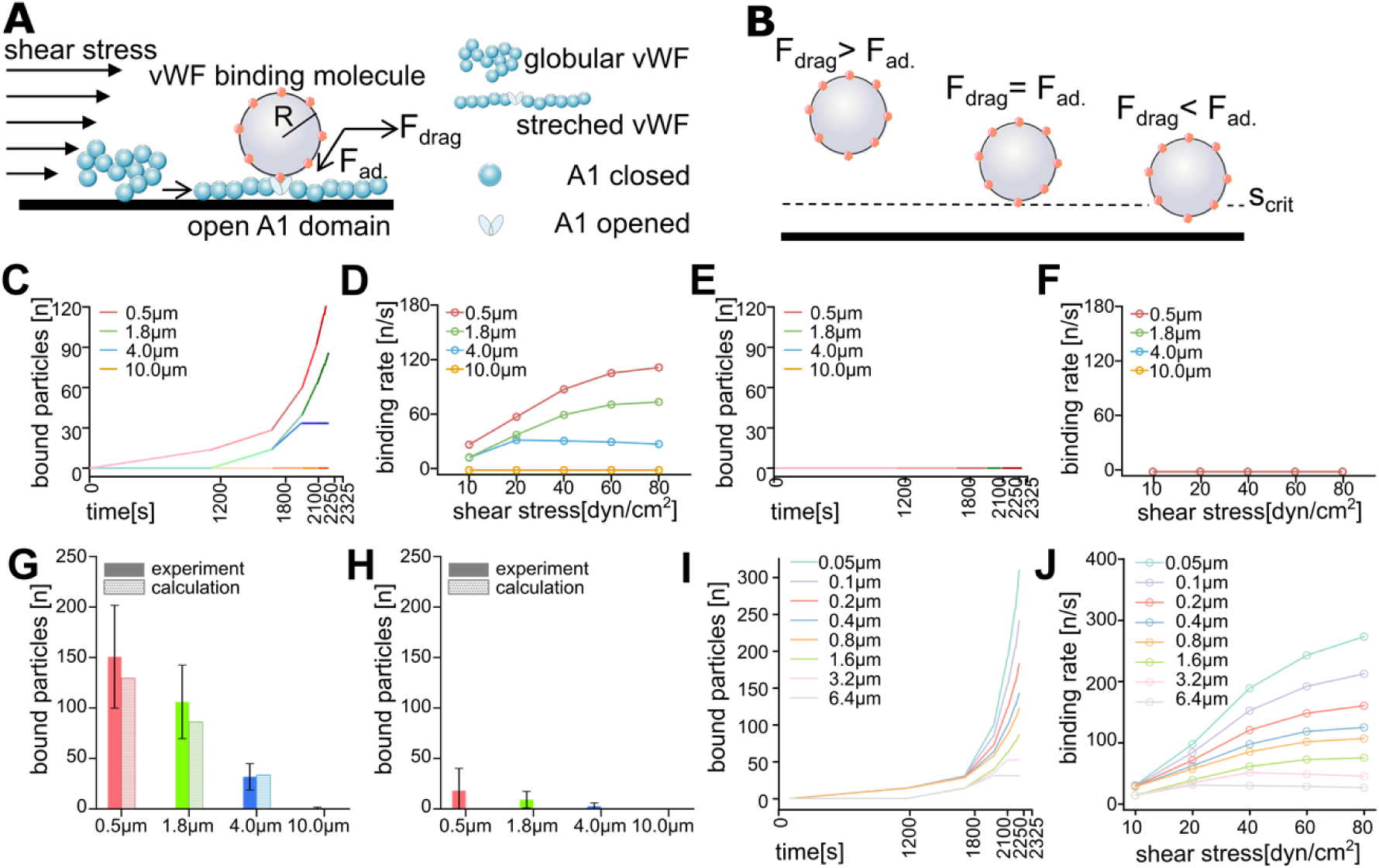
Theoretical calculation of particle binding to vWF under shear stress. (**A**) Schematic presentation of particle binding to vWF fibers under flow. As further outlined in the supplement, particle binding depends on the size of the particle (particle radius, R), the drag force (F_drag_) pulling at the particle and the adhesion force (F_ad_). Consistent with the experiment shown in Fig.2, we calculated the binding of particles to a WT vWF-coated surface as a function of time and shear stress. (**B**) The balance between F_drag_ and F_ad_ promote particle adhesion. At a separation above s_crit_, F_drag_ exceeds F_ad_ and particles detach. At a separation below s_crit_, F_drag_ is smaller than F_ad_ and particles bind. (**C**) Binding of particles to WT vWF-coated surface as a function of time and shear stress. (**D**) The particle binding rate as a function of shear stress was derived from the data shown in (C). (**E**) Binding of particles to ΔA1 vWF-coated surface as a function of time and shear stress. (**F**) The particle binding rate as a function of shear stress was derived from the data shown in (E). (**G**, **H**) Comparison of calculated and experimental results. Compared are the total number of particles bound to the WT vWF-(G) or ΔA1 vWF-(H) coated surfaces at the final time point. (**I**) Calculation of particle binding to WT vWF covering a wider size range of the particles. (**J**) Calculated binding rates of particles to WT vWF; related to the data shown in (I).

**Figure 4.**
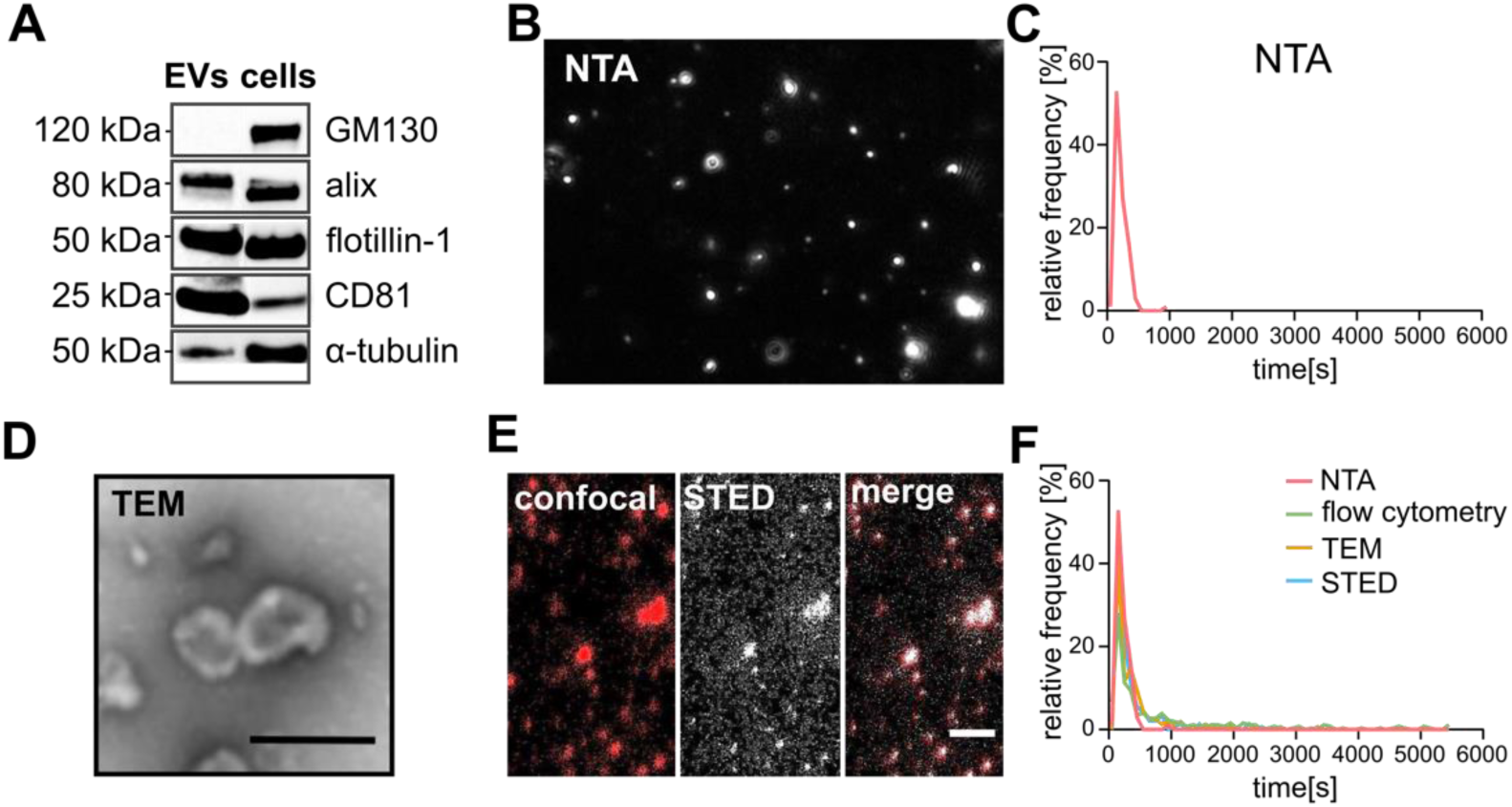
Characterization of B16F10 melanoma cell-derived EVs. (**A**) Western blot analysis of EVs and B16F10 cells. CD81 (22 kDa), flotillin-1 (48 kDa) and alix (96 kDa) were detected in EVs and cells. The Golgi resident protein GM130 served as a non-EV marker and was only detected in cells, α-Tubulin served as a loading control. (**B**) Representative NTA image and determined size distribution of EVs. (**C**) The particle diameter ranged from 118.8 nm to 324.9 nm, based on the 10th to 90th percentile NTA measurements. The mean diameter of the EVs was 213.9 ± 4.5 nm. (**D**) Representative transmission electron microscopy (TEM) image. Scale bar = 250 nm. (**E**) Confocal laser scanning microscopy (CLSM) image (red), STED image (white) and merge of both channels to indicate super-resolution of the STED. EVs were labeled with the lipophilic tracer DiD. Scale bar = 2 μm. (**F**) Size distributions of EVs obtained by NTA, flow cytometry, TEM and STED microscopy.

Consistent with the experiment shown in **Figure 2F** and **G**, none of the particles bound to the surface. For a better comparison of the experimental data (**Figure 2**) and the theoretical calculations (**Figure 3**), we compared the number of bound particles at the final time point. **Figure 3G** shows the comparison of the WT vWF coated surfaces; **Figure 3H** the comparison of the ΔA1 vWF coated surfaces. Experimental data and the theoretical calculations are in good agreement and strongly suggest that vWF under shear is a size-selective molecule able to trap small sized objects while exhibiting no capacity to bind larger ones.

Our experiments showed that particles with a diameter of 4 μm interact only weakly with vWF (**Figure 2D**) suggesting that 4 μm is close to a size threshold. In order to further pin down such a threshold, we have extended our calculation to include a wider range of particles (**Figure 3I**). While the adhesion of particles with a diameter of 1.6 μm still slightly increase even at higher shear stress, we noted that the adhesion rate of larger particles (3.2 μm and 6.4 μm) started to decrease at shear stresses above 40 or 20 dyn/cm², respectively (**Figure 3J**). Taken together, our data suggests that the size critical for adhesion is between 1.6 μm and 3.2 μm. It is interesting to note that the diameter of platelets, which are the best known binding partners of vWF, is within this critical size range. This highlights that the size of platelets seems to be an optimal compromise, integrating shear stress-dependent adhesion with a maximum cargo load capacity.

### 4. Characterization of melanoma cell-derived EVs

In the following, we aimed to transfer the finding that vWF is size-selective and able to bind small but not larger particles to a more physiological context. Since our results suggested that small-sized particles have an increased ability to bind to stretched vWF, we assume that EVs, rather than intact tumor cells, are capable of binding to vWF under shear stress. To validate our hypothesis, we isolated and characterized EVs from B16F10 metastatic melanoma cells, since these cells produced abundant EVs enriched in the HS-exposing proteoglycan glypican.^22, 33^

After isolation of EVs by ultracentrifugation, Western blot analysis was used for EV characterization using previously established marker proteins. Whole cell lysates served as a reference (**Figure 4A**). Alix (96 kDa) and the membrane-bound flotillin-1 (48 kDa) were present in the isolated EVs and the cell lysate. The Golgi resident protein GM130 (used as a marker for non-EV contamination) was detected only in cell lysates. In contrast, we found enrichment of the CD81 tetraspanin in EVs compared to cell lysates. Taken together these data indicate a high purity of our isolated EVs. Next, we measured the size distribution and concentration of the EVs. **Figure 4B** shows the results obtained by nanoparticle tracking analysis (NTA). The concentration of EVs was 1.61×10^9^ ± 3.74×10^7^ particles/mL. The mean diameter was 213.9 ± 4.5 nm, while EVs with a diameter of 165 nm were most abundant (**Figure 4C**). The detection range of NTA is limited to the size range between 50 nm and 1000 nm^34^, therefore we measured the size of the EVs by flow cytometry, transmission electron microscopy (TEM) (**Figure 4D**) and super-resolution stimulated emission depletion (STED) microscopy (**Figure 4E**). EV size distributions from all different methods were very consistent indicating that the majority of the EVs had a diameter of 118.8 nm to 324.9 nm (**Figure 4F**). Therefore, B16F10- derived EVs are below the critical size range of vWF and we assume that their ability to interact with vWF under shear is high. In contrast, melanoma cells have a diameter above 10 μm and we assumed a very low binding probability.

### 5. EVs but not whole cells bind to stretched vWF

To test whether EVs but not whole cells bind to vWF fibers, we perfused vWF coated surfaces with B16F10 cell-derived EVs, B16F10 cells, platelets and erythrocytes (45% haematocrit) (**Figure 5A**). A representative snapshot of the experiment (**Figure 5B**) shows that platelets form a pearl-on-string configuration indicating their interaction with vWF fibers. In contrast, B16F10 cells did not attach to the surface in any of our experiments. Instead, we observed a significant accumulation of EVs in close proximity to platelets, following the pearl-on-string-like arrangement (**Figure 5B**). A direct comparison revealed that EVs bound to vWF fibers less efficiently than platelets (**Figure 5C**), although EVs are smaller than platelets—and thus according to our theoretical model should bind more efficiently This discrepancy occurs because (i) many individual EVs (∼100-200 nm) fall below the resolution limit of conventional fluorescence microscopy, and (ii) when EVs cluster in close proximity, they are optically resolved as single fluorescent spots - both leading to systematic underestimation of EV binding events. Additionally, and as further outlined in the discussion, platelets are much softer than EVs and deform under shear stress.^35–38^ This deformation, which was not considered in our calculation, increases the area of adhesion and in turn promotes the interaction of platelets with vWF.

**Figure 5.**
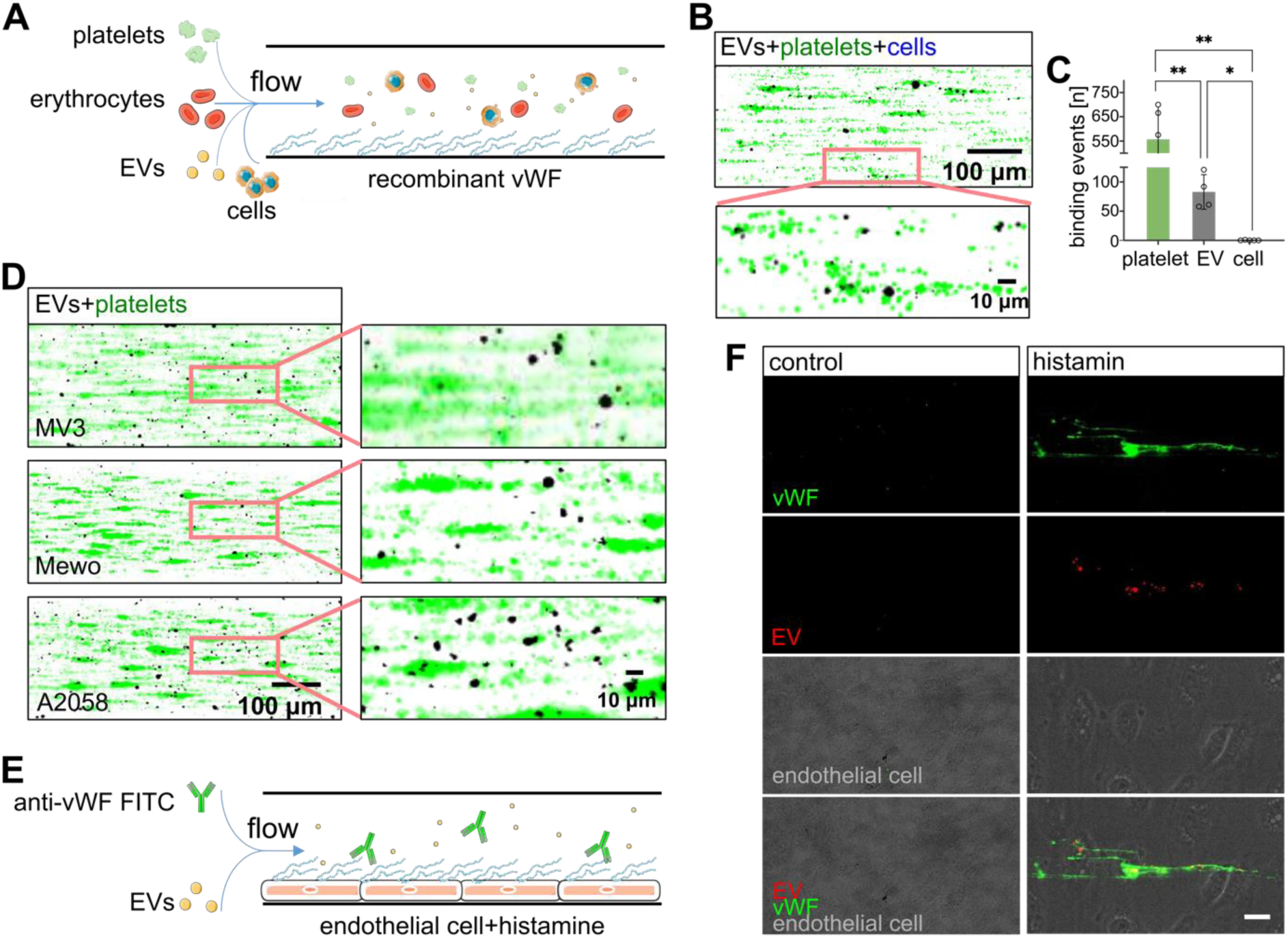
Binding of melanoma cell derived-EVs to stretched vWF under flow. (**A**) Schematic presentation of the microfluidic experiment. Melanoma cells and melanoma cell-derived EVs were perfused through vWF-coated channels in the presence of platelets and erythrocytes. (**B**) EVs (black) and platelets (green) bind to the vWF-coated surface. Pearl-on-string-like arrangement indicate the interaction of platelets and EVs with vWF fibers. Bound B16F10 cells (principally labelled in blue) were not detected. (**C**) Quantification of bound platelets, EVs and cells (n= 5). ** P ≤ 0.01 one-way ANOVA with Tukey post hoc test. (**D**) EVs (black) isolated from different melanoma cell lines (MV3, Mewo, A2058) bound to the vWF-coated surfaces. (**E**) Schematic presentation of the microfluidic experiment. HUVECs were coated on the bottom of channels, after their stimulation with histamine, vWF was released. EVs and FITC-conjugated antibodies directed against vWF were perfused through flow channels. (**F**) After histamine stimulation and the release of vWF, EVs (red) bound directly to vWF fibers (green). No EVs bound directly to the HUVECs (bright field). Scale bar = 20 μm. (Schematic elements used from Servier Medical Art: https://smart.servier.com/).

EVs were produced by most mammalian cells. To determine whether the interaction of EVs with vWF fibers is a general phenomenon and not restricted to EVs from B16F10 cells, we tested the binding of EVs from the human melanoma cell lines MV3, MeWo and A2058 to coated vWF. As shown in **Figure 5D**, EVs from all the different cell lines accumulate in a pearl on string configuration together with platelets. To rule out the possibility that EV binding is influenced by platelets, we repeated the experiment with B16F10 cell-derived EVs without the addition of platelets. Binding was still observed, which could be reduced by adding the vWF-cleaving protease ADAMTS13 (**S-Figure 5**).

In our initial approach, the evidence of the EV-vWF interaction was only indirect and indicated by platelet binding. To demonstrate direct and specific EV binding to vWF under physiological conditions, we perfused human umbilical vein endothelial cells (HUVECs) B16F10 cell-derived EVs (**Figure 5E)**. The release of vWF from the endothelium was induced by histamine stimulation. We stained the vWF fibers during the flow experiment with a fluorophore-conjugated antibody, which was added into the perfusing medium. After stimulation of HUVECs with histamine, EVs bound directly to the released vWF fibers, whereas no EV binding was detectable on non-histamine stimulated HUVECs (**Figure 5F**).

### 6. EVs binding to vWF depends on the A1 domain and HS

Our previous studies suggested that HS at the surface of cells is a relevant ligand for the A1 domain of vWF. ^1^ To prove whether HS is also involved in the binding of EVs to vWF, we isolated EVs from B16F10 cells lacking the HS-synthesizing enzyme exostosin 1 (EXT1, B16F10 *Ext1-/-* cells). EXT1 is a key enzyme of the HS biosynthesis and its knockout completely prevent HS expression^1^. As shown in **Figure 6A**, we perfused vWF-coated surfaces either with EVs derived from WT cells or *Ext1-/-* cells together with platelets and erythrocytes. To study the shear stress dependent binding of EVs, we performed again force-ramp experiment (**Figure 6B**). In comparison to EVs isolated from WT cells, HS-deficient EVs isolated from *Ext1-/-* cells bound less to the vWF-coated surface (**Figure 6C**). Consistent with our previous experiments, the number of bound EVs increased over the course of the experiment and with increasing shear stress for both types of EVs (**Figure 6D**). However, the binding rate of the HS-deficient EVs was strongly decreased, although the overall shear stress dependent binding profile was similar to WT cell-derived EVs (**Figure 6E**). Quantitative evaluation at the endpoint of multiple experiments indicated that HS deficiently EVs bound more than two times less efficient to vWF than the EVs from WT cells (**Figure 6F**).

**Figure 6.**
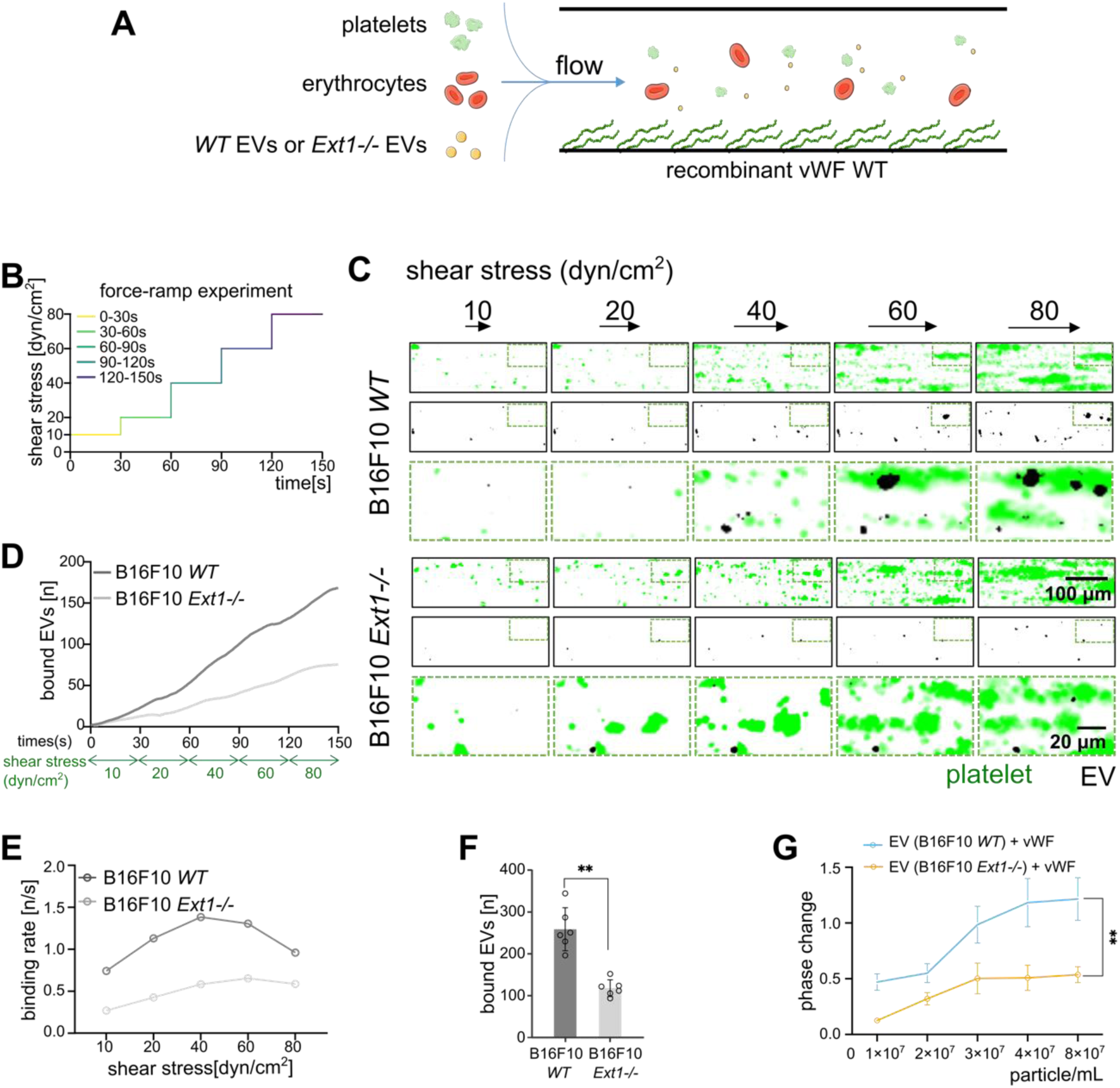
Binding of melanoma cell derived EVs to stretched vWF depends on HS. (**A**) Schematic representation of the microfluidic experiment. EVs derived from WT and *Ext1-/-* cells were perfused through microfluidic channels coated with recombinant WT vWF in the presence of platelets and erythrocytes. (**B**) Force-ramp experiments were performed with a time-dependent change of shear stress. (**C**) Time-lapse dual-color fluorescence images of the interaction of platelets (green) and EVs (black) from B16F10 WT or *Ext1-/-* cells with vWF at the indicated shear stress. The merged images showed a zoom in on the region indicated by the dashed box. (**D**) Binding of B16F10 WT or *Ext1-/-* EVs (black) to vWF as a function of time and shear stress. (**E**) Binding rates of B16F10 WT or *Ext1-/-* EVs to vWF-coated surfaces indicated that EV binding increases with increasing shear stress until 40 dyn/cm² was reached. (**F**) Quantification of bound EVs at the end of multiple experiments (n=7). ** P ≤ 0.01 Student’s t test. (**G**) SAW biosensors were coated with WT vWF followed by the addition of B16F10 WT or *Ext1-/-* EVs at different concentrations. The increase in phase shift indicate the binding of EVs. Compared to *Ext1-/-* EVs, the steeper increase of the phase shift after the addition of WT EVs indicate a significant (P ≤ 0.01 Student’s t test) stronger binding affinity to vWF (n=3). (Schematic elements used from Servier Medical Art: https://smart.servier.com/).

Our data, shown in **Figure 2**, suggested that PE-particles interact with the A1 domain of vWF. As depicted in **S-Figure 6A-E**, lack of the A1 domain reduced also the EV binding rate and the total number of bound particles. Similarly, the presence of the vWF cleaving protease ADAMTS13 prevented the accumulation of EVs at the channel surface (**S-Figure 6F-H**). Taken together, the experiments shown in the **Figures 6**, **S-Figure 5** and **S-Figure 7** suggest that stretched vWF is able to bind EVs and that HS is the ligand for the A1 domain. As an internal standard, we always added platelets into the perfusing medium. In the presence of ADAMTS13, the binding of platelets was not affected at shear stress below 60 dyn/cm². Above 60 dyn/cm², platelet binding was clearly reduced by ADAMTS13 **(S-Figure 6I, J).** The cleavage site of ADAMTS13 is located within the A2 domain of vWF, which, like the A1 domain, is cryptic in globular vWF but accessible to ADAMTS13 when vWF is stretched. We noted the same trend of a reduced platelet binding on surfaces coated with the ΔA1 vWF (**S-Figure 6I, J**); however, the binding of platelets was not completely abolished on ΔA1 vWF coated surfaces. Although this requires further investigation, it may indicate that platelet-derived WT vWF contributes to the adhesion process and at least partially compensates for the lack of the A1 domain. As expected, platelet binding was not different between channels perfused with EVs isolated from HS-deficient cells or wild-type cells (**S-Figure 6I, J**).

**Figure 7.**
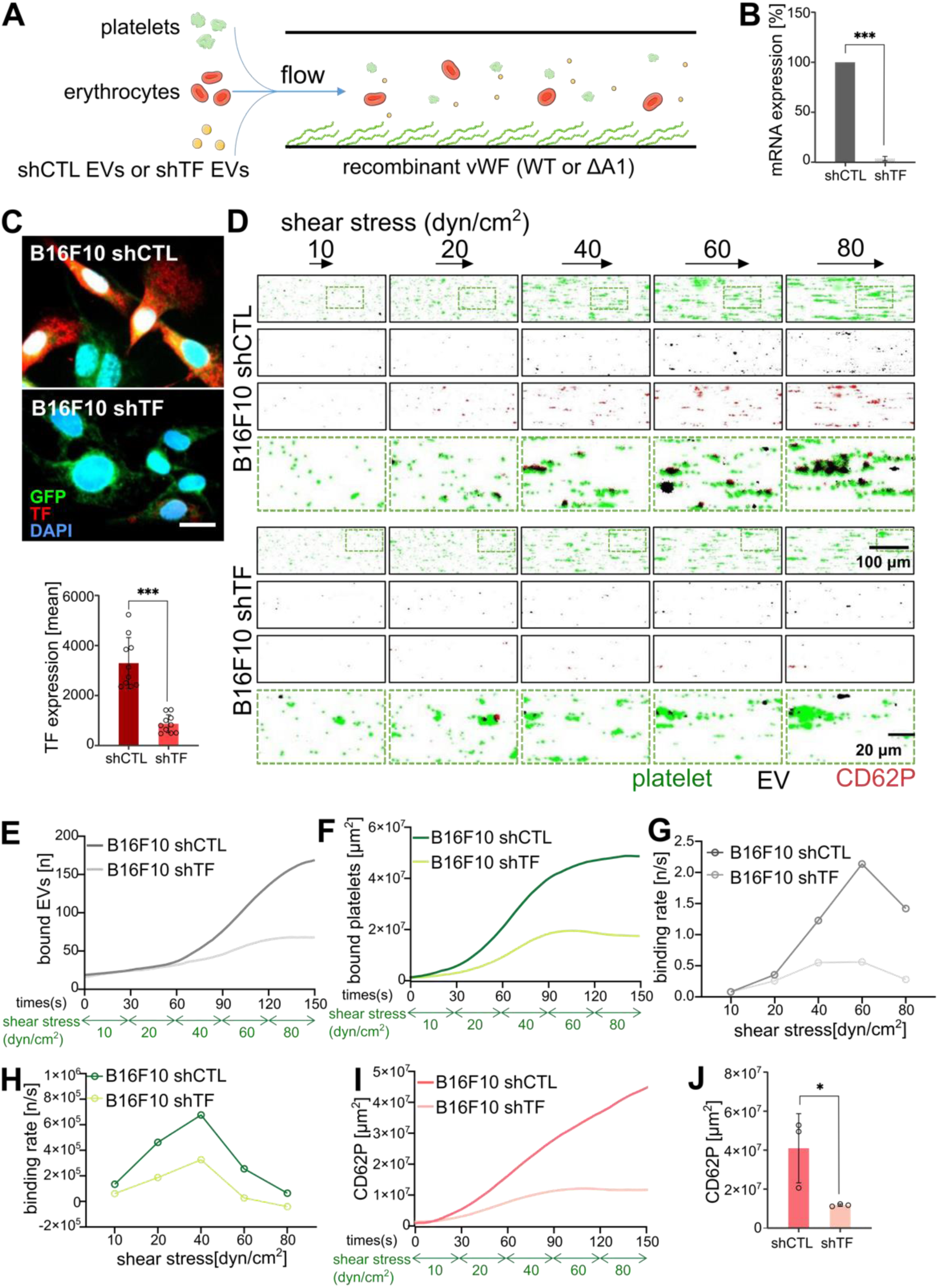
Binding of EVs to vWF induces platelet activation. (**A**) Schematic representation of the microfluidic experiment. EVs were perfused in microfluidic channels coated with recombinant vWF in the presence of platelets, plasma and erythrocytes. (**B**) Significantly decreased TF expression in B16F10 shTF compared to shCTL cells was measured by qPCR (n = 3) and (**C**) immunofluorescence staining (n = 10). Scale bar = 20μm. ** P ≤ 0.01 Student’s T test. (**D**) Time-lapse dual-color fluorescence images show the binding of platelets (green) and shCTL or shTF-EVs (black) to vWF at the indicated shear stresses. CD62P was used as marker for activated platelets (red). The merged images showed a zoom in on the region indicated by the dashed box. (**E**) Binding of shCTL and shTF EVs to vWF-coated surfaces as a function of shear stress and time. (**F**) Binding of platelets to vWF-coated surfaces in the presence of shCTL or shTF EVs as a function of shear stress and time. (**G**) Binding rate of shCTL and shTF EVs to vWF-coated surfaces as a function of shear stress. Data were derived from (E). (**H**) Binding rates of platelets to vWF-coated surfaces in the presence of shCTL or shTF EVs as a function of shear stress. Data were derived from (F). (**I**) Platelet activation during the flow experiment as a function of shear stress and time. Activation was compared in channels containing shCTL EVs or shTF EVs. (**J**) Quantification of EV induced platelet activation at the end of the flow experiment (n=3). * P ≤ 0.05, *** P ≤ 0.001 Student’s t test. (Schematic elements used from Servier Medical Art: https://smart.servier.com/).

To further assess the specificity and strength of the EV-vWF interaction, we employed a surface acoustic wave (SAW) biosensor assay. This label-free biosensing approach allowed us to quantify the real-time binding kinetics of B16F10 *WT* and *Ext1-/-* EVs to WT vWF-coated surfaces. Upon the addition of WT EVs, a pronounced and dose-dependent phase shift was observed, indicating strong binding to vWF (**Figure 6G**). In contrast, *Ext1-/-* EVs induced a significantly lower phase shift, reflecting a markedly reduced binding affinity. Parallel experiments with ΔA1 vWF-coated surfaces revealed diminished phase shifts for WT EV types (**S-Figure 7A**), demonstrating that A1-domain deletion substantially reduced their binding.

### 7. Binding of EVs to vWF fibers support platelet aggregation

Due to the simultaneous binding of EVs and platelets along the vWF fibers, TF, which is frequently exposed on EVs, is positioned in close proximity to the platelets. In the presence of plasma, TF produces thrombin, a strong activator of platelets^39^. We adjusted our microfluidic setup to study the potential impact of TF on platelet activation and the shear dependent binding of EVs and platelets (**Figure 7A**). Consistent with our previous experiments, we separated erythrocytes and platelets from fresh blood and performed force ramp experiments.

Additionally, we added plasma together with CaCl_2_ into the perfusing medium to allow coagulation. The plasma obtained from healthy volunteers contained physiological levels of ADAMTS13. To prevent the cleavage of vWF fibers and to mimic a pathophysiological lack of ADAMTS13, channel surfaces were coated with vWF lacking the A2 domain (ΔA2 vWF). The A2 domain harbors the cleavage site for ADAMTS13 and thus, ΔA2 vWF is not cleavable in the presence of plasma. We compared the effect of EVs derived from TF knockdown cells (B16F10 shTF) and EVs derived from control cells (B16F10 shCTL). The shRNA-induced knockdown of TF was quantified by qRT-PCR and immune fluorescence staining (**Figure 7B, C**).

**Figure 7D** shows representative images taken during the microfluidic experiment at different shear stresses. Consistent with our previous experiments, increasing shear stress increased the binding of EVs (**Figure 7E**) and platelets (**Figure 7F**). However, calculating the corresponding binding rates revealed that the critical shear stress for EV and platelet binding was lower compared to experiments without plasma (**Figure 7G, H**). In our previous experiments (**Figures 1** and **6**), performed in the absence of plasma, the critical shear stress was 60 dyn/cm²; now, in the presence of plasma, we measured a critical shear stress of 40 dyn/cm². This agrees well with the larger platelet aggregates and the resulting increase in drag force at higher shear flow. The shift of the critical shear stress was more pronounced in channels perfused with the EVs from B16F10 shCTL cells than in channels containing the EVs from B16F10 shTF cells. Consistent with that, also the size of the platelet aggregates grew steeper in the control cell group (**Figure 7D**). To better prove that platelets were activated, we added a fluorescence-conjugated antibody directed against P-selectin (CD62p) to the perfusing medium. As shown in **Figure 7D and Video S2**, platelet exposed increasing amounts of P-selectin at the surface in the course of the experiment. P-selectin presentation was more prominent in channels containing the EVs from the shCTL cells (**Figure 7I, J**). Platelet activation and the formation of larger platelet aggregates indicate blood clotting within the channels resembling the situation of microvessels in patients with hypercoagulation. ^39–41^

To further confirm the impact of TF, we compared EVs from the melanoma cell lines B16F10 and A2058, which express different levels of TF.^4^ TF expression levels were quantified using fluorescence microscopy, flow cytometry, and Western blot **(S-Figure 8A–C)**. The characterization of A2058-derived EVs is shown in **S-Figure 8D**. We performed microfluidic experiments with EVs derived from B16F10 cells in the presence of CaCl₂ and plasma. **S-Figure 8E** illustrates the binding of EVs to stretched vWF under increasing shear stress. The binding of EVs from both cell lines was similar **(S-Figure 8F, G)**. Additionally, the area covered by bound platelets reached comparable levels regardless of the EV type added. However, in the presence of B16F10-derived EVs, the platelet-covered area increased more rapidly at lower shear stress compared to the group treated with A2058-derived EVs **(S-Figure 8H, I)**. Moreover, platelet aggregation was significantly higher in the presence of B16F10-derived EVs **(S-Figure 8J, K)**. Taken together, our data suggest that EVs bound to vWF promote platelet activation and aggregation. In the following, we intend to further understand whether the interplay of EVs, vWF, platelet and the plasmatic coagulation is able to trap circulating tumor cells.

**Figure 8.**
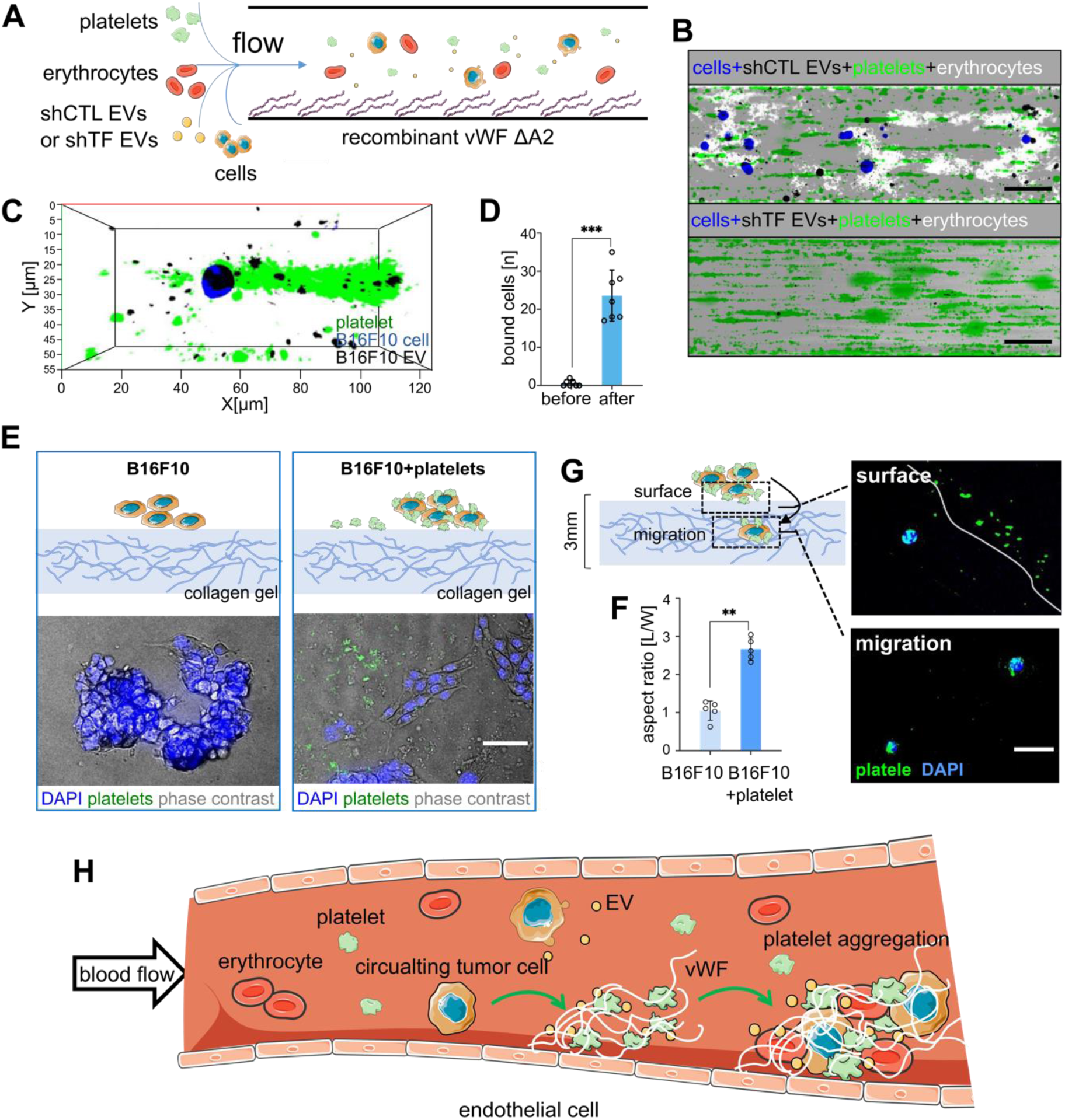
Conglomerates of EVs, platelets and vWF trap tumor cells. (**A**) Schematic presentation of the microfluidic experiment. B16F10 cells and EVs either isolated from chCTL or shTF cells were perfused through vWF-coated channels in the presence of platelets, plasma and erythrocytes. (**B**) B16F10 cells (blue) bound to vWF in channels containing EVs isolated from shCTL B16F10 cells. Binding of the melanoma cells was associated with the aggregation of erythrocytes (white). No B16F10 cells or erythrocytes accumulated in channels perfused with EVs isolated from shTF B16F10 cells. Scale bar = 100μm. (**C**) 3D rendered image showing the interaction of a melanoma cell (blue) with platelets (green) and EVs (black). The image was generated from optical sections acquired by structured illumination microscopy. (**D**) Numbers of trapped B16F10 cells before or after EV-platelet conglomerates are formed (n=7). *** P ≤ 0.001 Student’s t test. (**E**) B16F10 cells were seeded on collagen gels with or without thrombin activated platelets. In the absence of platelets, melanoma cells were round shaped and formed clusters. In the presence of platelets, melanoma cells showed an elongated shape and no clustering. Scale bar = 50 μm. (**F**) Quantification of the aspect ratio of B16F10 cells in the presence or absence of activated platelets. ** P ≤ 0.01 Student’s T test. (**G**) Migration of B16F10 cells incubated with platelets from the surface to the center of the collagen gel. Representative fluorescence images show the gel surface (surface of the gel is indicated by the white line) and the central part of the gel (migration). Platelets (green) accumulated at the gel surface and around single melanoma cells (blue). Scale bar = 50 μm. (**I**) Schematic summary of the presented data. Under shear flow, vWF released from activated endothelial cells is stretched and binds platelets as well as EVs. EVs can be released from circulating tumor cells or other blood cells such as platelets. In the presence of plasmatic coagulation factors, the binding of TF-exposing EVs promotes platelet activation and aggregation. Although tumor cells do not directly bind to stretched vWF, platelet aggregation and thrombus formation subsequently entraps erythrocytes and circulating tumor cells. (Schematic elements used from Servier Medical Art: https://smart.servier.com/).

### 8. Tumor cells are trapped in platelet aggregates

Based on the experimental setup shown in **Figure 7**, we added wild-type B16F10 melanoma cells into the perfusing medium (**Figure 8A**). While our data, shown in **Figure 5**, indicate that melanoma cells do not directly bind to stretched vWF, we now found that melanoma cells are trapped within large thrombus-like aggregates composed of platelets, EVs and erythrocytes (**Figure 8B, C**). The quantitative evaluation, presented in **Figure 8D**, shows that EV induced platelet aggregation increased significantly the number of trapped tumor cells.

Previous studies have indicated that platelets support the dissemination of tumor cells and potentially their escape from the blood vessel^42^. To follow the effect of the entrapment of B16F10 cells in aggregates of activated platelets, we seeded B16F10 cell on a soft collagen matrix with or without activated platelets. After an incubation time of 48 h the morphology and migration of the cells were measured by fluorescence microscopy (**Figure 8E**). In the absence of platelets, B16F10 cells formed clusters and the cells had a round morphology that is characteristic for non-adherent and less viable cells. In contrast, B16F10 cells interacting with activated platelets displayed an elongated morphology, indicating adherent and viable cells. To quantify the different cell morphologies, we measured the length and the width of the cells and calculated the aspect ratio. **Figure 8F** shows that platelet free cells had an aspect ratio of 1.1 ± 0.10, whereas platelet-associated cells had a significantly increased ration of 2.7 ± 0.15. In addition, only B16F10 cells associated with platelets were able to migrate into the collagen gel (**Figure 8G**). During their migration, tumor cells escaped from the clot but kept a shell of platelets, which was detectable even in deeper regions of the collagen matrix.

Taken together, our data indicate that vWF is a size-selective protein that favors the binding of small sized objects such as EVs. Once co-trapped with platelets, EVs promote their activation and the formation of larger clots that could incorporate circulating tumor cells. The results of our study are summarized schematically in **Figure 8H**.

## Discussion

The findings presented in this study shed light on the intricate interplay between vWF fibers, EVs and cells. In the past, many molecules and cells have been identified as ligands for vWF indicating that vWF is an important protein within our circulation and involved in a large range of biological processes. Multimeric vWF is considered as key player in coagulation but may also contribute to other endothelial functions such as angiogenesis^43^ or immune cell recruitment^13, 44^. In the present study, we investigated the balance between the shear force-dependent opening of the A1 domain promoting adhesion and the shear force-dependent drag force counteracting adhesion. In line with previous studies, opening of the A1 domain started in our microfluidic experiments at a shear stress of around 20 dyn/cm². ^3, 7, 45^ To measure the impact of the drag force in a standardized way, we utilized PE particles with a size range from 0.5 to 10 μm. We found that smaller particles (<4 μm) have a higher ability to interact with vWF coated surfaces. Larger particles (≥ 4 μm) in the size of cells were unable to attach, suggesting that the drag force exceeded the adhesion force. Similar results were obtained in our calculations and also in experiments with whole melanoma cells and melanoma cells derived EVs. Taken together, our data show that vWF is a size-selective protein able to distinguish between small and large particles under flow. In future studies, we aim to extend our theoretical calculations by advanced computer simulations considering additional parameters such as the potential effect of particle margination or the deformation of cells under flow. ^46, 47^

In comparison to platelets, EVs bound less efficiently, although according to our calculation, their size would favor an even stronger binding ability. Our theoretical calculation considers simple sphere-shaped particles that cannot deform under shear stress. Platelets are however, soft structures with a comparable low elastic modulus^35, 36^ promoting their flattening under shear stress. Flattening increases the adhesion area of platelets and thus their net ability to adhere to a vWF-coated surface. EVs are much smaller and their elastic modulus was found to be relatively high. ^48^ In comparison to platelets, it can therefore be assumed that the ability of EVs to flatten under flow is low and that in turn the adhesion area is not affected by shear stress.

Not only platelets but also larger cells such as leukocytes or tumor cells may have the ability to deform under flow, which may increase the area of adhesion. Previous studies already indicate that tumor cells with a lower elastic modulus are more metastatic suggesting also an increased ability to attach to the vascular wall. However, the extent to which vWF fibers directly contribute to vascular metastasis required further investigation^23^. In the present study, our data suggests that not vWF fibers directly but rather conglomerates of vWF, EVs and platelets are required to trap circulating tumor cells as a potential starting point for tumor cell extravasation and metastasis.

The formation of vWF-platelet strings is a locally restricted process that enables the response of a defined vascular area to endothelial cell activation or injury. The binding of platelets to vWF via GPIbα is a transient process and contraction of the vWF fiber leads to the dissociation of the platelets. ^49^ Therefore, the formation of stable aggregates requires an additional activation of platelets and a stabilization of the vWF-platelet conglomerate through fibrin. Thrombin, the downstream product of TF, is a key enzyme facilitating the activation of platelets through protease-activated receptor signaling and the conversion of fibrinogen into fibrin. ^50^ Protease-activated receptor signaling in platelets triggers the exposure of GPIIb/IIIa, which binds to the RGD motif of vWF. ^50, 51^ After its local production, thrombin is rapidly diluted in the circulation and due to a relatively short life-time eliminated within seconds. ^52^ In the present study, we found that EVs bind to vWF fibers in close proximity to platelets. This increases the likelihood that thrombin, after being produced by TF on the surface of EVs, will be lossless transferred to the nearby platelets. The prometastatic role of platelets and TF-induced coagulation is well documented and the direct physical interaction of platelets and tumor cells is known to protect the tumor cells against the attack of the immune system and to support metastasis. ^8, 24, 53, 54^ This is consistent with our data, showing the improved migration of B16F10 cells through collagen gels in the presence of EV-vWF-platelet conglomerates.

Our experiments indicate that tumor cell-derived EVs bound to vWF via HS, which is ubiquitously expressed by mammalian cells, suggesting that vWF fibers might also capture EVs from other cell types. Given that EVs are produced by almost all types of cells, their levels in the blood circulation are high. While most EVs originate from platelets, those derived from leukocytes, red blood cells, and circulating tumor cells also contribute to blood coagulation. ^17^ Interestingly, previous studies discovered that EVs derived from sickle red blood cells or neutrophils promote the release of vWF from the endothelium to enhance thrombotic processes and future experiments will show whether the binding of EVs to vWF fibers contribute to thrombosis in such diseases. ^55–57^

It has been reported that the amount of TF^+^-EVs in the blood increases under various disease conditions. ^27, 58^ Therefore, we hypothesize that the binding of TF^+^-EVs to vWF is not limited to cancer-associated thrombosis, but is a general mechanism contributing to vWF-mediated platelet aggregation and coagulation, e.g. to seal vascular injuries or to contribute to hypercoagulation in disease. EVs were postulated as biomarkers predicting disease progression and severity. For example, clinical observations suggested that vWF together with EVs from gastric cancer cells are associated with cancer aggression and poor clinical outcomes for patients. ^59^ Also, the exposure of TF by EVs was shown to be linked to lung metastasis in a murine tumor model. ^24^ Here, we found that lack of HS decreased the adhesion of EVs to vWF fibers and previously we have shown that lack of HS on melanoma cells is connected to metastasis. ^1^ Whether lack of HS is related to reduced EV-mediated thrombosis requires further research, but may offer a therapeutic targeting e.g. by blocking the HS binding site of vWF by HS mimetics.

## Conclusions

In conclusion, our study shows that the binding of EVs to stretched vWF could activate platelets and trigger platelet aggregation. These clots have the ability to trap circulating tumor cells suggesting a connection between the binding of EVs to vWF and metastasis. By elucidating the molecular and biophysical mechanisms underlying EV-vWF interactions, we offer a foundation for future therapeutic interventions targeting EV-mediated thrombosis in cancer patients and in other diseases characterized by hypercoagulation. Additionally, our findings underscore the clinical relevance of EVs and vWF as potential biomarkers for assessing thrombotic risk and monitoring disease progression. In our study, we focused on the crosstalk between EVs and vWF fibers; however, the impact of blood rheology on the interplay between blood flowing EVs and other components of the vascular system such as leukocytes or endothelial cells, has not yet been investigated in detail. Therefore, future research is required to fully understand the impact of blood flow on the biological effects of EVs in health and disease.

## List of abbreviations

ADAMTS13: A disintegrin and metalloproteinase with a thrombospondin type 1 motif, member 13
AFM: Atomic force microscope
ΔA1: deletion of the A1 domain
ΔA2: deletion of the A2 domain
EVs: Extracellular vesicles
EXT1: Exostosin 1
F_ad_: Adhesion Force
F_drag_: Drag Force
GPIbα: Glycoprotein Ib alpha
HUVECs: Human umbilical vein endothelial cells
HS: Heparan sulfate/heparin
NTA: Nanoparticle tracking analysis
PE: Polystyrene
SMFS: Single molecule force spectroscopy
STED: Stimulated emission depletion
TEM: Transmission electron microscopy
TF: Tissue factor
vWF: von Willebrand factor

## Declarations

### Ethics approval and consent to participate

All animal experiments were approved by the governmental animal care authorities (“Behörde für Justiz und Verbraucherschutz, Hamburg” project N033/2022).

### Consent for publication

All authors have agreed to publish this manuscript.

### Availability of data and materials

All data generated or analysed during this study are included in this published article and the Supplementary materials. For original data, please contact corresponding author.

### Competing interests

The authors have no competing interests related to this work.

### Funding

This study was supported by research funding from the Mildred Scheel Cancer Career Center HaTriCS4 at University Medical Center Hamburg-Eppendorf, the German Research Foundation within priority program 2416 CodeChi GO 2528/10-1, or in terms of GRK 2873 *‘Tools and Drugs of the Future’* and the Hamburg Pro Exzellenzia Plus Scholarship Program.

### Authors’ contributions

Y.W., X.L., and C.G. designed the study and wrote the manuscript. Y.W., X.L., T.D., A.T., K.N., and S.B. performed experiments and subsequent analyses. Y.W., X.L., A.T.B., B.P., A.T., G.B., D.F. and D.A.F. analyzed and discussed data. C.G. and S.W.S. edited the manuscript.

X.L. and Y.W. share the first author position based on their equal contributions for the study.

All authors reviewed the manuscript.

## Acknowledgments

The authors would like to thank Dr. A.V. Failla from the UKE Microscopy Imaging Facility for his assistance with the STED microscope. We are grateful to Sabine Vidal-y-Sy and Tobias Obser for their excellent technical support. Special thanks go to Prof. Dr. Francisco M. Goycoolea (Goycoolea Cell Biology and Histology, Faculty of Biology, University of Murcia, Murcia, Spain) for generously providing the HS-coated fluorescent nanocapsules used in our mouse injection experiments.

## Notes

### Competing Interest Statement

The authors have declared no competing interest.

