## supplementary material for "Binding of extracellular vesicles to stretched von Willebrand factor promotes platelet activation"

Running title: Size-selective binding of EVs to stretched vWF

Yuanyuan Wang[1,2+], Xiaobo Liu[1+], Tomasz Downar[3], Alper Topuz[4], Alexander T. Bauer[1], Katrin Nekipelov[5], Santra Brenna[6], Gerd Bendas[5], Berta Puig[6], Stefan W. Schneider[1], Dmitry A. Fedosov[4], Christian Gorzelanny[1]

**Supplementary information:** calculation of particle binding under shear stress.

**Supplementary Figures 1-8**

- **Supplement Figure 1:** Flow rate response to shear stress in microfluidic system.
- **Supplement Figure 2:** Binding of heparin coated PE particles to vWF.
- **Supplement Figure 3:** Binding of blood flowing objects to vWF under shear stress.
- **Supplement Figure 4:** Analysis of the HS-vWF Interaction by single molecular force spectroscopy.
- **Supplement Figure 5:** Binding of EVs to vWF in the absence of platelets.
- **Supplement Figure 6:** Binding of melanoma cell-derived EVs to stretched vWF mutants.
- **Supplement Figure 7:** Binding of EVs to vWF mutant measured by SAW measurements.
- **Supplement Figure 8:** Binding of EVs from different different TF-expressing cell lines to vWF.

**Supplementary videos 1-2**

**Supplementary information**

To validate our experimental data shown in Figure 1, we calculated the binding of PE-particles to vWF at different shear stress conditions as illustrated schematically in S-Figure 1. We calculated the drag force ($F_{drag}$) pulling at the particle by Stoke's law:

$$F_{drag} = 6\pi R \mu \gamma (R + s_0) \qquad (1)$$

where μ is the viscosity of the perfusing medium (0.01 dyn s/cm$^2$), γ the shear stress and $s_0$ the separation of the particle from the surface. R is the radius of the particle. We considered the drag force only of particles that are very close to the vascular wall where $R >> s_0$.Equation 1 is therefore simplified to:

$$F_{drag} = 6\pi R^2 \mu \gamma \qquad (2)$$

To estimate the adhesion force ($F_{ad}$), we used the previous approach of Jiang and co-workers, which have calculate the interaction force of GPIbalpha and the A1 domain of vWF.[1] To estimate the interaction force of HS and vWF, we performed single molecular force spectroscopy using an atomic force microscope (S-Figure 2). We evaluated our data assuming that the rupture of the HS-vWF interaction is related to the applied loading rate as previously described[2]. We found a bond length of 0.399 ± 0.096 nm and an off-rate of 1.6 ± 0.8 1/s. Our data are in good agreement with previous measurements of the interaction of different vWF A1 domain constructs with GPIbα, which documented a bond length range from 0.44 to 1.4 nm and off-rates between 3.27 and 10.8 1/s.[3] Therefore, we assumed that the binding of HS-coated particles is steered by the same positively charged amino acids of the A1 domain. The interaction force between the A1 domain and HS depends strongly on their separation, s, as indicated by equation 3:

$$F_{A1-HS}(s) = \sum_{i,j} -\frac{q_i q_j}{4\pi\varepsilon_0\varepsilon} \frac{e^{\kappa r}}{(1+\kappa r)} \frac{(\kappa s + 1)e^{-\kappa s}}{s^2} + \frac{6C_{vdw}}{s^7} \qquad (3)$$

where the charges, $q_i$, of the positively charged aminoacids of the A1 domain have been listed in the publication of Jiang et al.[1] The charges of HS, $q_j$, were set to 1. $\varepsilon$ and $\varepsilon_0$ are the relative and the vacuum dielectric constants, respectively. The radii, *r*, of the hydrated charged groups were estimated to be in the range of 3.5 Å and, at a monovalent salt concentration of 150mM, the debye length $1/\kappa$ was calculated to be 7.7 Å.

We added to the previously published equation[4] a term accounting for weak van der Waals interactions promoting particle adhesion at short distances and independent of the electrostatic interaction. The corresponding constant $C_{vdw}$ was set to -6.66x10$^{-83}$ J m$^6$.

Further, we assumed that multiple A1 domains can interact with distinct binding partners at the particle surface. The distance, *d*, of the binding partners was estimated to be 500nm and the resulting separation s' of the multiple binding sites was calculated using equation 4:

$$s' = \cos\left(\tan^{-1}\frac{d}{R+s_0}\right)\left(\sqrt{(R+s_0)^2 + d^2} - R\right) \qquad (4)$$

where $s_0$ accounts for the shortest distance between the adhering particle and the surface as indicated in the schematic drawing shown in S-Figure 1A. To estimate the adhesion force ($F_{ad}$) in three dimensions, we extended equation 2:

$$F_{ad}(s) = \left[\frac{2\pi R}{d}\sum_{l} F_{A1-HS}(s')\right] + F_{A1-HS}(s_0) \tag{5}$$

We aimed to calculate the number of particles bound to a vWF-coated surface at different shear stress conditions. Increasing shear stress promotes the elongation of vWF and opening of the A1 domain. The percentage of open A1 domains at defined shear stress was quantified by measuring the adhesion of platelets to vWF. At a wall shear stress of 4 dyn/cm², 1.5% of the A1 domains were open, at 8 dyn/cm² 5.2%, at 16 dyn/cm² 12.5%, at 32 dyn/cm² 52%, at 64 dyn/cm² 80%, and at 128 dyn/cm² 98%. Additionally, increasing shear stress is directly linked to an increased flow rate and therefore to an increased amount of particles flowing over the adhesive surface. The number of particles, $n$, accumulating over time in the field of view (FoV) of the microscope is linked to the wall shear rate $\dot{\gamma}_w$:

$$n(t) = \frac{w\dot{\gamma}_w h^2}{2}\, tc_{total} \tag{6}$$

where $w$ is the width of the FoV, $h$ is the height of the FoV, $t$ is the time of the experiment and $c_{total}$ the total particle concentration ($3 \times 10^{12}$ particels/ml). The wall shear stress is linked to $\dot{\gamma}_w$ through the viscosity, $\mu$.To estimate the number of particles close enough to the vWF-coated surface to be trapped, we defined a critical separation, $s_{crit}$, of the flowing particles from the surface. As illustrated in S-Figure 1B, $s_{crit}$ defines the separation when $F_{drag}$ is equal to $F_{ad}$. To link $s_{crit}$ to the number of bound particles, $n_{bound}$, we adjusted equation 6:

$$n_{bound}(t) = \frac{w\dot{\gamma}_w s_{crit}^2}{2}\, tc_{total} \tag{7}$$

Note that in our calculation we have ignored the potential effect of particle margination toward the wall.[5, 6] We expect that particle margination will not significantly change the results, since particle migration toward the wall is slow and is likely not very pronounced in relatively short microfluidic channels used in the experiments.

### Supplementary Figures

#### Figure S1, related to Figure 1

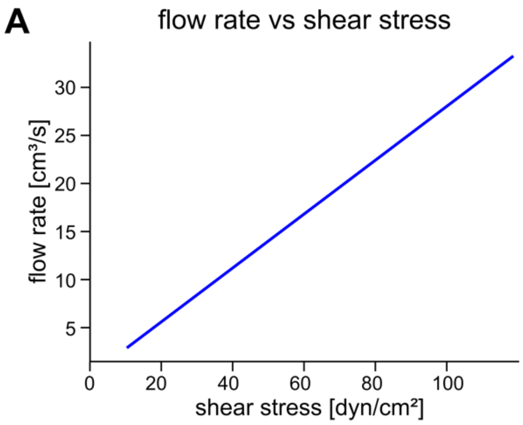


**Supplement Figure 1. Flow rate response to shear stress in microfluidic system. (A)** Shown is the relationship between the flow rate and the shear stress , demonstrating monotonic increase from 0 to 30 $cm^3$/s across the tested range (0-80 $dyn/cm^2$).

### Figure S2, related to Figure 2

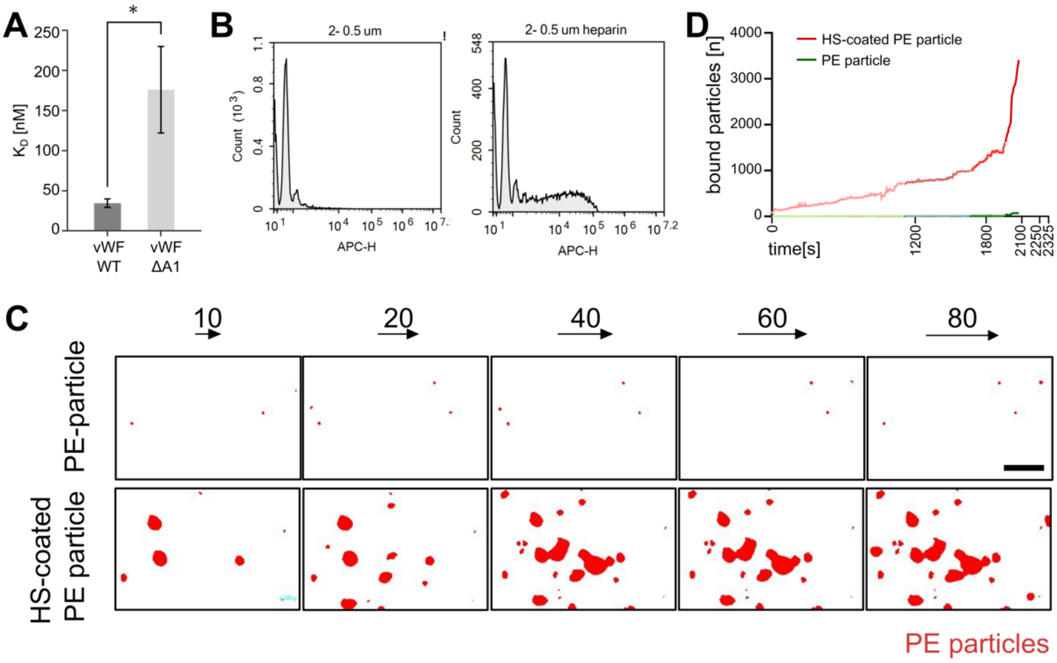


**Supplement Figure 2. Binding of heparin coated PE particles to vWF.** (**A**) Dissociation contant ($K_D$) of the interaction between heparin with WT or ΔA1 vWF (n=3). (**B**) Heparin coating efficiency on PE-particles was measured by flow cytometry. The rightward shift of the peak indicates successful heparin conjugation to the particle surface. (**C**) Representative microscopy images of non-coated (upper) and heparin-coated (lower) PE-particls (red) bound to vWF-coated surface at the specified shear stress. Scale bar =10 µm. (**D**) Binding of particles to vWF under time-correlated shear stress conditions: The color gradient indicate the change of the shear stress from 10 to 80 dyn/cm². * P ≤ 0.05 Student's t test.

**Figure S3, related to Figure 3**

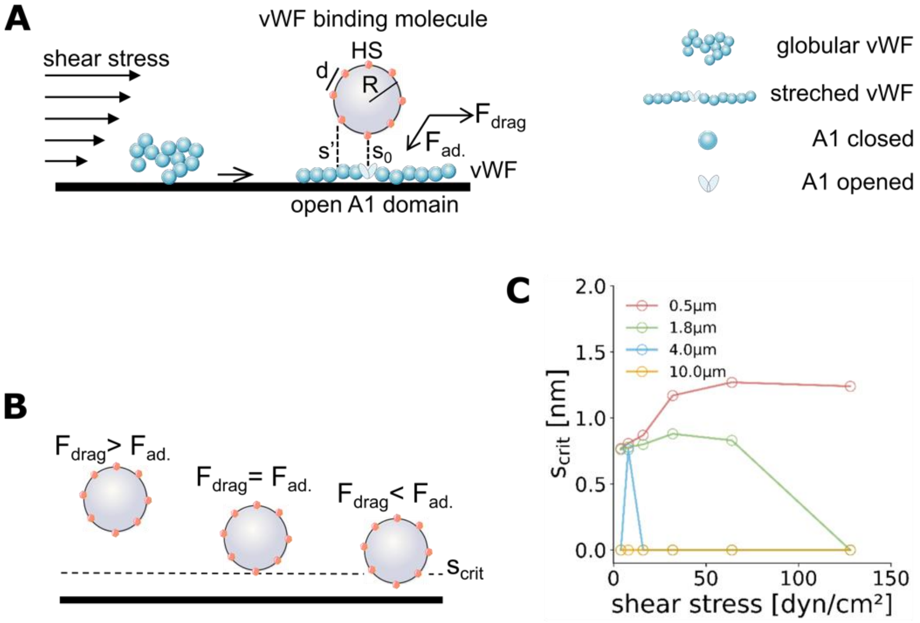


**Supplement Figure 3. Binding of blood flowing objects to vWF under shear stress.** (**A**) Schematic drawing illustrating the interaction between HS-coated particles and vWF under shear flow. The drag force ($F_{drag}$) acting on a particle is determined by Stokes' law, where R is the particle radius, s0 and s' are the separations between the particle and the surface at different positions. The adhesion force ($F_{ad}$) is a result of the interaction between vWF and HS. (**B**) Schematic presentation how the balance between $F_{drag}$ and $F_{ad}$ promote particle adhesion. At a separation above $s_{crit}$, $F_{drag}$ exceeds $F_{ad}$ and particles detach. At a separation below scrit, $F_{drag}$ is smaller than $F_{ad}$ and particles bind.(**C**) Shown is the change of $s_{crit}$ as a function of the applied shear stress and for particles with different diameters (0.5µm, 1.8µm, 4.0µm, 10.0µm).

**Figure S4, related to Figure 3**

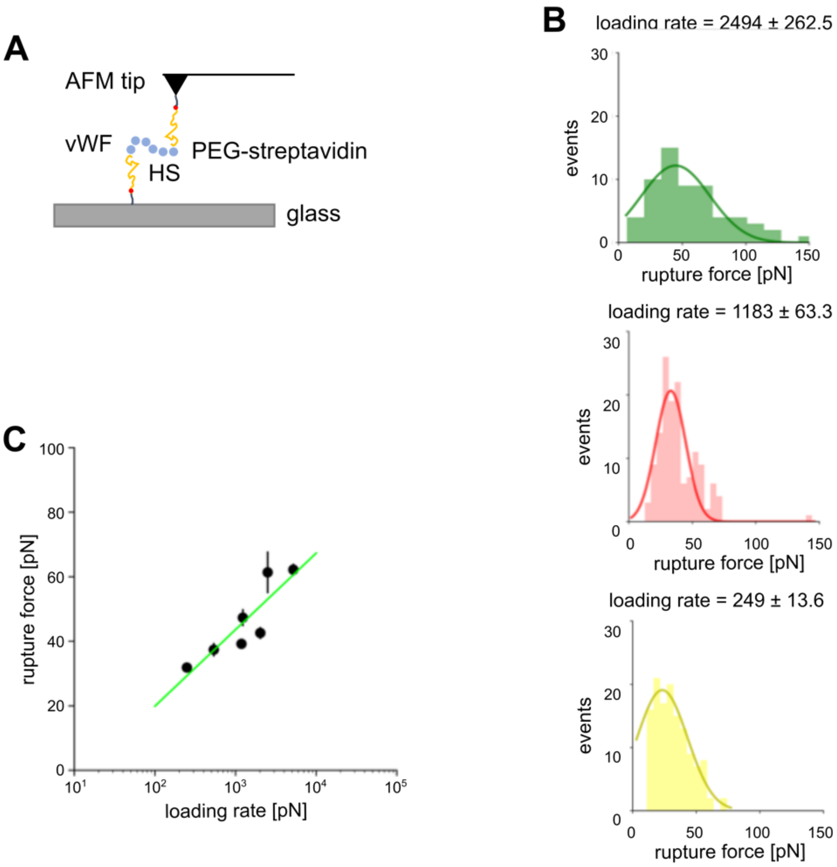


**Supplement Figure 4. Analysis of the HS-vWF Interaction by single molecular force spectroscopy.** (**A**) Schematic representation of the conducted single-molecule force spectroscopy experiment. Biotinylated HS was attached to the tip of the atomic force microscope and the glass substrate through polyethylene (PEG)-linker immobilized streptavidin. Recombinant vWF was added to the glass slide shortly prior to the measurement. (**B**) Distribution of rupture forces at indicated loading rates. The histograms illustrate the probability distribution of rupture forces required to separate HS from vWF. (C) Relationship between rupture force and loading rate.

**Figure S5, related to Figure 5**

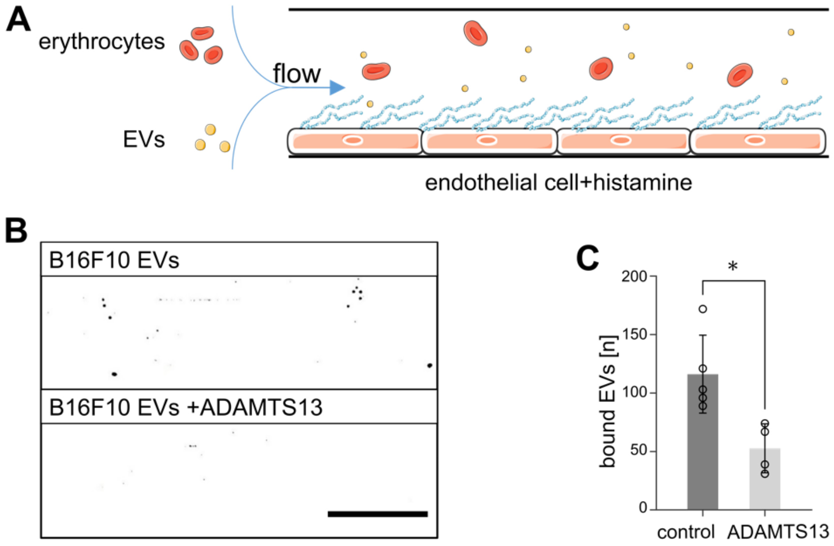


**Supplement Figure 5. Binding of EVs to vWF in the absence of platelets.** (**A**) Schematic presentation of the microfluidic experiment. EVs were perfused through channels coated with recombinant vWF and erythrocytes. (**B**) Representative images of EV binding to vWF with or without ADAMTS13 treatment. (**C**) Quantification of EV binding to vWF (n=4). * $P \leq 0.05$ Student's t test.

**Figure S6, related to Figure 6**

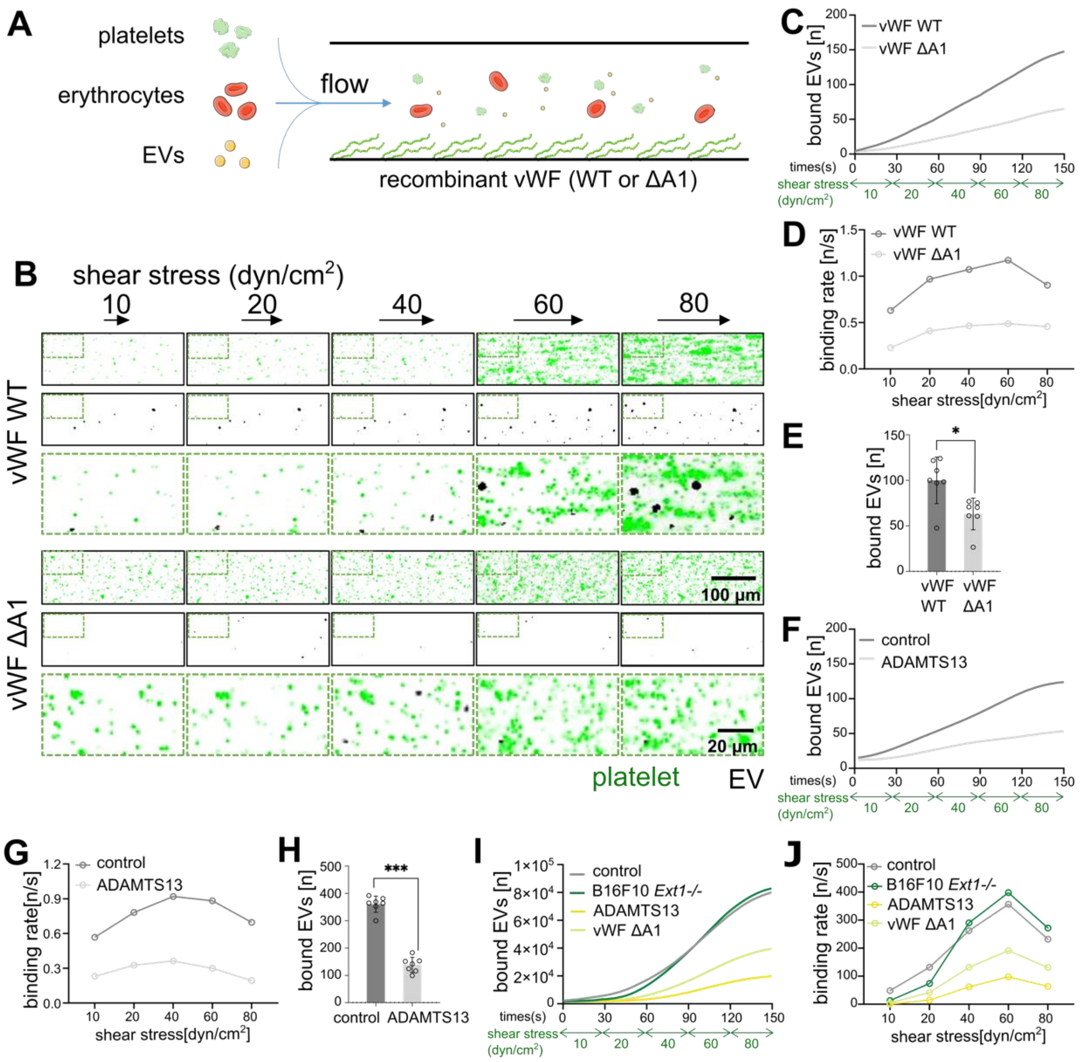


**Supplement Figure 6. Binding of melanoma cell-derived EVs to stretched vWF mutants.** (**A**) Schematic presentation of the microfluidic experiment. EVs were perfused through channels coated with recombinant WT or ΔA1 vWF. (**B**) Time-lapse dual-color fluorescence images of the interaction of platelets (green) and EVs (black) with either WT vWF or ΔA1 vWF under different shear stress conditions. The merged images showed a zoom in on the region indicated by the dashed box. (**C**) Binding of EVs to WT or ΔA1 vWF as a funtion of time and shear stress. (**D**) Binding rate of EVs to WT or ΔA1 vWF as a funtion of shear stress. (**E**) Number of bound EVs at the end of the experiment (n=7). (**F**) Binding of EVs to WT vWF in the presence or absence of ADAMTS13 as a funtion of time and shear stress. (**G**) Binding rate of EVs to WT vWF in teh presence or absence of ADAMTS13 as a funtion of shear stress. (**H**) Number of bound EVs at the end of the experiment (n=7). Data are presented as the mean ± SD, *** P≤ 0.0005 Student's t test. (**I**) Comparative analysis of the binding of platelets to vWF (WT or ΔA1) in the presence or absence of ADAMTS13 as a function of shear stress and time. (**J**) Binding rate of platelets to vWF (WT or ΔA1) in the presence or absence of ADAMTS13 as a function of shear stress. Generated from data shown in (I). Data are presented as the mean ± SD, * P≤ 0.05 Student's t test. *** P≤ 0.001 Student's t test.

**Figure S7, related to Figure 6**

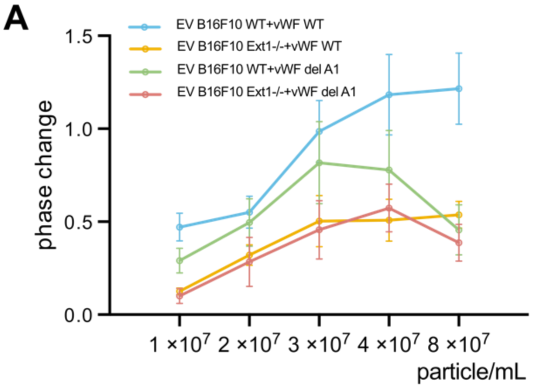


**Supplement Figure 7. Binding of EVs to vWF mutant measured by SAW measurements.** (A) Quantification Phase change after the addition of B16F10 WT or Ext1-/- EVs to ΔA1 vWF reflecting binding kinetics and interaction strength. (n=3)

**Figure S8, related to Figure 7**

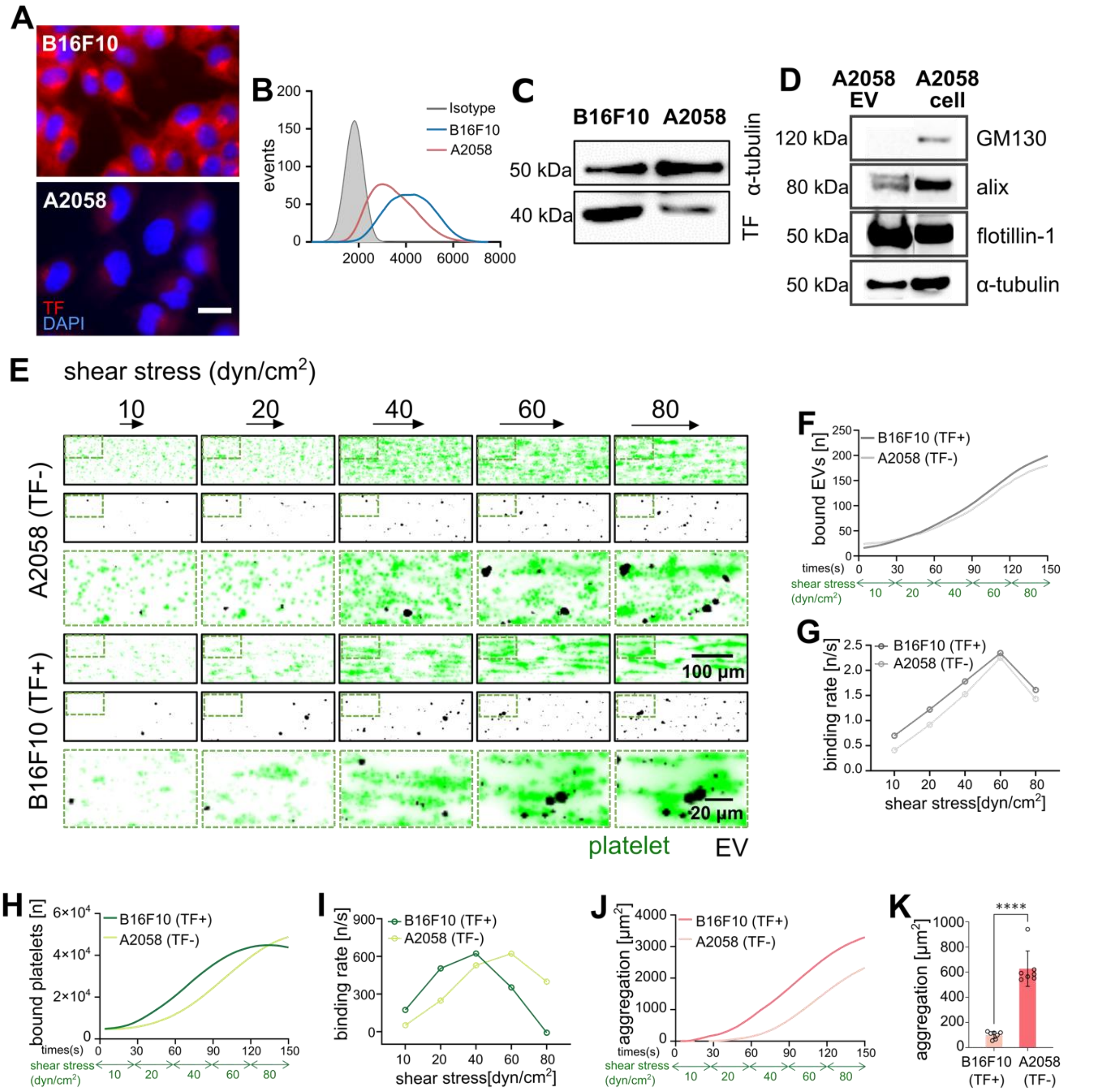


**Supplement Figure 8. Binding of EVs of different TF-expressing cell lines to vWF.** (**A**) Immunofluorescence staining of TF (red) on B16F10 and A2058 melanoma cells. Nuclei are stained by DAPI (blue). Scale bars = 20µm (**B**) Quantification of TF expression on B16F10 and A2058 cells by flow cytometry and (**C**) by Western blot. α-Tubulin served as a loading control. (**D**) Western blot analysis of EV isolated from A2058 cells. CD81 (22 kDa), flotillin-1 (48 kDa) and alix (96 kDa) were detected in the EVs and cell lysate. The Golgi resident protein GM130 served as a non-EV marker, α-Tubulin served as a loading control. (**E**) Binding of platelets (green) and EVs (black) isolated from A2058 or B16F10 cells with vWF at the indicated shear stresses. The merged images showed a zoom in on the region indicated by the dashed box. (**F**) Binding of EVs to vWF coated surfaces as a funtion of time and shear stress. (**G**) Binding rate of EVs to vWF-coated surfaces as a funtion of shear stress. (**H**) Binding of platelets to vWF coated surfaces in the presence of EVs isolated from B16F10 cor A2058 cells as a funtion of time and shear stress. (**I**) Binding rate of EVs to vWF-coated surfaces as a funtion of shear stress. (**J**) Platelets aggregation in the presence of EVs as a funtion of shear stress and time. (**K**) Quantification of EV induced platelet aggregation at the end of the flow experiment (n=7). Data are presented as the mean ± SD, **** P≤ 0.0001 Student's t test.

**Supplementary videos**

**Video S1, related to Figure 1**

**Video S2, related to Figure 7**
